# Dynamic changes to the plastoglobule lipidome and proteome in heat-stressed maize

**DOI:** 10.1101/2025.06.13.659543

**Authors:** Elsinraju Devadasu, Febri A. Susanto, Anthony L. Schilmiller, Cassandra Johnny, Peter K. Lundquist

## Abstract

Heat stress is a major environmental factor affecting the physiology and productivity of agricultural crops including maize (*Zea mays*). Plastoglobules, lipid-protein structures in chloroplasts, play a key role in stress resilience by modulating lipid metabolism and maintaining chloroplast function. However, the molecular functions of plastoglobules and their compositions are enigmatic. Our study investigated the molecular changes in the protein and lipid compositions of plastoglobules and thylakoids at six time points over the course of an imposed heat stress and recovery treatment in B73 inbred maize. Results indicate a progressive increase in plastoglobule size and number, and proliferation of adjacent cytosolic lipid droplets, correlating with the duration of heat exposure. Significant alterations in lipid composition, particularly in levels of triacylglycerol, plastoquinone derivatives (PQ-C, B & E) and the fatty acid phytol ester, 12:0-phytol, suggest a protective role in membrane remodeling and oxidative defense. Furthermore, heat-induced upregulation of key plastoglobule-associated proteins, such as Fibrillin 1a & 2, Fructose-bisphosphate Aldolase 2, 13-Lipoxygenase 10/11, and Allene Oxide Synthase 2b were observed, indicating their involvement in stress mitigation. These findings provide novel insights into the adaptive mechanisms of plastoglobules under heat stress in the context of remodeling at the thylakoid and highlight potential targets for improving maize resilience and leveraging the plastoglobules for crop improvement. Understanding these responses could contribute to developing heat-resilient maize cultivars in the face of global climate change.

## INTRODUCTION

Rising global temperatures present a profound challenge to global maize (*Zea mays*) productivity (Alexander et al., 2006; Lobell et al., 2011; Lobell et al., 2013; Sillmann et al., 2013). The associated heat stress interferes with numerous processes within the maize plant, negatively impacting growth, development and reproduction. These negative consequences are the result of a complex interplay of molecular and cellular activities. For example, prolonged elevated temperatures can promote precocious developmental transitions, ultimately limiting end-of-season yields (Tebaldi and Lobell, 2018). However, many of the detrimental effects from heat stress emerge even from short exposure durations of a few hours to a few days. Increases in temperature rapidly affect membrane fluidity and protein structure, rippling through the essential processes of the cell. Thus, heat waves, regardless of the duration, are harmful to crop productivity and require rapid cellular responses to mitigate negative effects.

The chloroplasts represent a key component in the sensing of – and adaptation to – heat stress (Teige, 2019; Hu et al., 2020; Zeng et al., 2021). The impact of heat stress is felt acutely at the photosynthetically active thylakoid membrane where a delicate balance of supply and demand of photosynthates must be maintained to prevent spill-over of excited electrons out of the transport chain that can quickly lead to uncontrolled oxidative damage. The thylakoid lipids are composed primarily of the galactolipids, monogalactosyldiacylglycerol (MGDG) and digalactosyldiacylglycerol (DGDG), along with more minor amounts of the sulfolipid, sulfoquinovosyldiaclyglycerol (SQDG), and phospholipid, phosphatidylglycerol (PG). The acyl groups of the galactolipids are comprised predominantly of the 18:3 polyunsaturated fatty acid, α-linolenic acid (Cook et al., 2021). However, under elevated temperatures which increase membrane fluidity, remodeling of the membrane acyl composition is quickly induced including turnover of 18:3 acyl groups through lipase activity. In *A. thaliana*, Heat Inducible Lipase 1 (AtHIL1) has been found to be partially responsible for the removal of 18:3 from MGDG, and a T-DNA knock-out mutant, *hil1*, is compromised in tolerance to short-term heat stress, underlining the importance of thylakoid acyl remodeling to proper heat stress adaptation (Higashi et al., 2018). Homologs of HIL1 are found in most land plants, including maize, and maintain similar co-expression networks, indicating that the role of HIL1 in 18:3 removal of MGDG under heat stress is widely conserved among plants.

Oxidation of membrane lipids and proteins is a significant aspect of heat-induced damage. Components of the photosynthetic machinery are sensitive targets, particularly the photosystem II complex (PSII). Direct damage to the photosynthetic machinery, including denaturation and side chain oxidation, results in photoinhibition and acceleration of the PSII repair cycle. Likewise, the prevalent polyunsaturated 18:3 acyl groups of the membrane lipids are readily peroxidized through reaction with hydroxyl radicals or hydrogen peroxide. The resulting peroxidized lipid radicals then enter a chain reaction generating amplified levels of peroxide radical formation among lipids, leading to membrane structural damage. Antioxidants, such as α-tocopherol, are necessary to quench (*i.e.*, terminate) the chain reaction. Additionally, enzymatic pathways can divert peroxided lipids into various oxylipins such as green leaf volatiles (via Hydroperoxide Lyase), or the phytohormone jasmonic acid (JA; via Allene Oxide Synthase) which hold stress-signaling capacity (Mosblech et al., 2009; Okazaki and Saito, 2014; Borrego and Kolomiets, 2016; Savchenko et al., 2017; Wasternack and Feussner, 2018; Lee and Kim, 2024).

To prevent or mitigate the generation of ROS arising from over-reduction of the electron transport chain, multiple processes are initiated in plants. Among these is non-photochemical quenching (NPQ) which describes the harmless dissipation of excited electrons as heat energy (Muller et al., 2001; Murchie and Ruban, 2020). Various reactive oxygen species (ROS)-scavenging methods are also employed including enzymatic antioxidant systems such as Superoxide Dismutase, Catalase, Ascorbate Peroxidase, and Glutathione Reductase, as well as non-enzymatic antioxidants such as carotenoids, ascorbic acid, tocopherols, and glutathione. These systems work in tandem to mitigate oxidative stress and maintain cellular stability. If left unchecked, ROS will react with lipids resulting in harmful membrane lipid peroxidation or the production of byproducts like β-cyclocitral, a lipid-derived volatile compound involved in stress adaptation in plants (Ramel et al., 2012; D’Alessandro et al., 2018; D’Alessandro and Havaux, 2019).

Plastoglobules, ubiquitous lipid-protein droplets of chloroplasts, are thought to play a prominent role in stress adaptation and membrane remodeling (Lundquist et al., 2012; Lundquist et al., 2020). Their physical connection to the thylakoid membrane and their dynamic morphological changes in response to numerous stresses, both biotic and abiotic, indicate that they facilitate remodeling of the thylakoid membrane through rapid and carefully regulated exchange of lipids and proteins between the two sub-compartments. Significantly, plastoglobules are conserved across the photosynthetic lineage. In fact, plastoglobule-like lipid droplets of *Synechocystis* sp. PCC6803 have recently been characterized, revealing shared features and a likely common evolutionary relationship. This suggests a critical role - or roles - in the support of photosynthetic life.

The investigation of *A. thaliana* has identified a unique plastoglobule proteome comprising approximately 35 proteins with exclusive, or nearly exclusive, plastoglobule localization. Due to the monolayer leaflet nature of the surface, and hydrophobic interior of plastoglobules (Austin et al., 2006; Lundquist et al., 2020), associated proteins are presumed to localize at the plastoglobule surface in a peripheral membrane manner. Many of the proteins of the *A. thaliana* proteome are known or putative enzymes involved in lipid and/or redox metabolism, including enzymes of the JA biosynthetic pathway, tocopherol biosynthesis, carotenoid catabolism, and lipid redox metabolism (Vidi et al., 2006; Ytterberg et al., 2006; Lundquist et al., 2012). A working model of plastoglobule function has been put forth in which plastoglobules serve as a metabolic platform to orchestrate various processes associated with thylakoid membrane remodeling (Lundquist et al., 2012; Lundquist et al., 2020). This model anticipates targeted changes to the proteome or lipidome in response to stress. The nature of these changes may vary depending on the nature and magnitude of the stress. Indeed, comparative proteome and lipidome characterization of *A. thaliana* plastoglobules from unstressed and high light-stressed leaf tissue has identified specific changes associated with leaf senescence and JA biosynthesis including recruitment of Plastoglobular Metallopeptidase M48 (PGM48), Esterase/Lipase/Thioesterase family 4 (ELT4), and Allene Oxide Synthase (AOS) to the plastoglobule, as well as a shift of specific carotenoid species from thylakoid to plastoglobule (Espinoza-Corral et al., 2021).

While the lipid and protein composition of *A. thaliana* plastoglobules is now relatively well understood, their composition in other species, including crop plants, remains uninvestigated. This hinders our understanding of how plastoglobules may be leveraged for agricultural improvement. Furthermore, comparative studies of plastoglobules during stress remain limited to the investigation of unstressed and high light-stressed *A. thaliana* plastoglobules by Espinoza-Corral et al. (2021).

In this manuscript we describe our detailed investigation of isolated *Z. mays* plastoglobules and thylakoids from six time points during a heat stress and recovery time-course. Proliferation of plastoglobules is observed by transmission electron microscopy (TEM) under the stress treatment and includes the observation of a unique crescent staining pattern of many plastoglobules specifically in bundle sheath (BS) cells. Proliferation of plastoglobules coincides with targeted effects on the proteome including the recruitment of multiple enzymes of the Calvin-Benson cycle and JA biosynthesis. Lipidome profiling identifies triacylglycerol (TAG) species, numerous derivatives of plastoquinone-9 (PQ-9), and a single FAPE species with a short chain saturated acyl group. Changes to the plastoglobule proteome and lipidome during the heat stress time-course are distinct from that described under water-deficit stress suggesting that *Z. mays* plastoglobules are deployed in differing ways to adapt to these two abiotic stresses, possibly related to different manifestations of the stresses within the thylakoid membranes.

## RESULTS

### Physiological consequences of heat stress

A heat-stress treatment was applied to four-week-old B73 inbred *Z. mays* by increasing the ambient air temperature in the greenhouse from 25 °C to 30 °C for one day and then to 40 °C for another seven days (**Figure 1A, Supp Fig S1**). Plants were then returned to 25 °C growth temperatures for an additional 3 days of recovery. During the heat stress, plants were watered at more frequent intervals to limit water-deficit conditions in the plant. Leaf relative water content (RWC; %) did not decline below *ca.* 80% at any time during the treatment, indicating that plants faced no or minimal water-deficit stress (**Supp Fig S1A, Supp Table S1**). In contrast, leaf temperatures increased to 34 – 37 °C during the stress treatment from a pre-stress temperature of *ca.* 28 °C (**Supp Fig S1B and C**). The elevated temperatures coincided with the emergence of chlorosis and curling at leaf tips of the older leaves, as well as wilt of the older leaves (**Figure 1B & 1C**).

**Figure 1.**
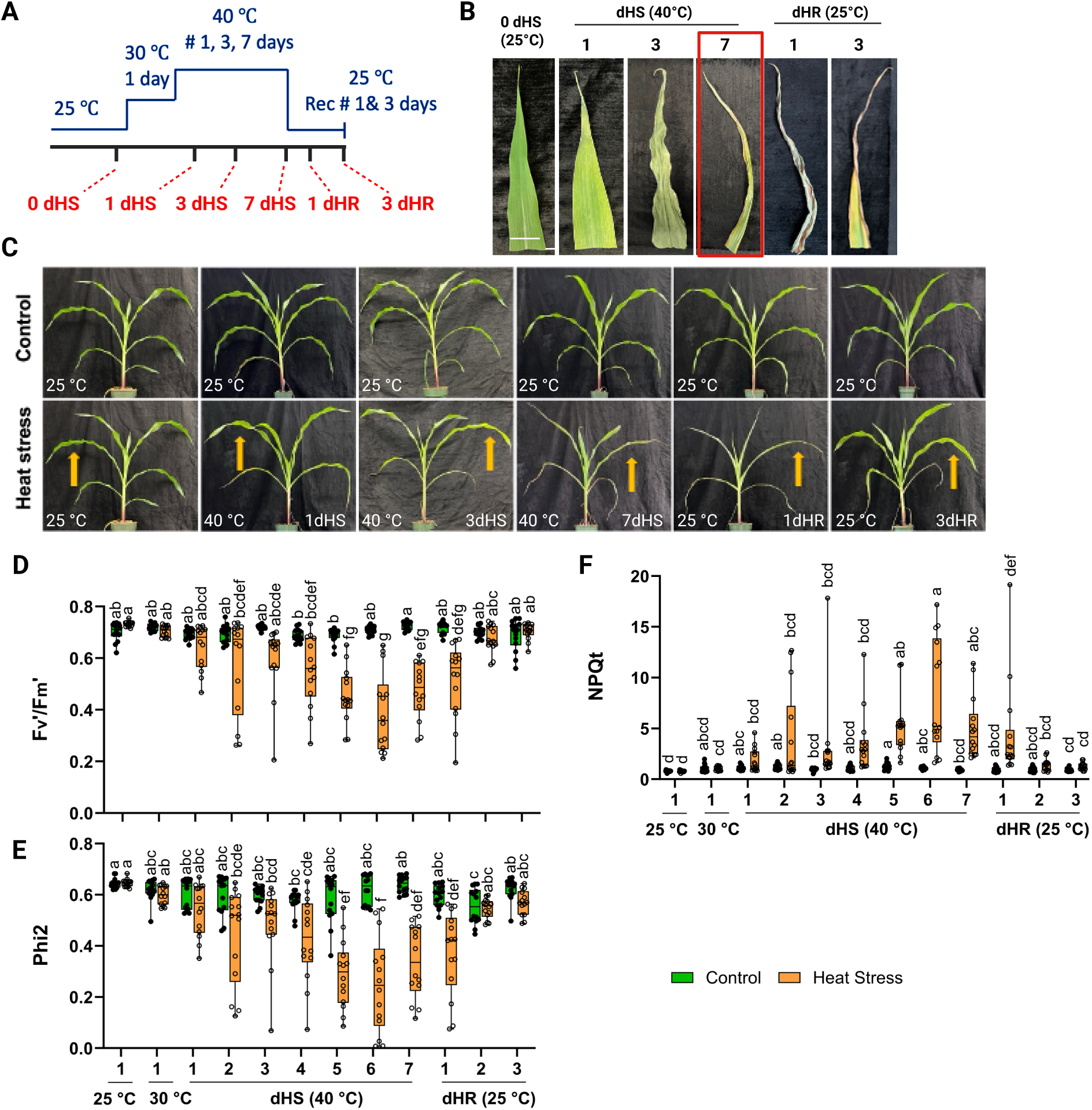
Phenotypic and photosynthetic measurements of maize leaves during a heat stress and recovery time-course. **A**) Illustration of the heat stress time-course imposed on B73 inbred maize. Time on the x-axis is not drawn to scale. Three-week-old maize, grown at 25 °C, were shifted to 30 °C for one day and then to 40 °C for another 7 days. Plants were then shifted back to 25 °C for a three-day recovery period. **B)** Representative photographs of leaf tips of the fifth collared leaf of maize plants at specified time points during the time course (white scale bar = 5 cm). The peak stress time point, 7 days of heat stress at 40 °C (7 dHS), is highlighted with a red box. **C)** Images in the top row are from control plants retained at 25 °C of growth temperature at the same time points, while those below represent the plants subjected to the heat stress. Yellow arrows indicate the fourth collared leaf used for photosynthetic measurements. **D-F)** Photosynthetic traits measured with a handheld MultiSpeQ chlorophyll fluorometer of the fourth collared leaf. Box and whisker plots represent the 25th and 75th quartiles with individual data points overlaid as open circles. n = 12 biological replicates (*i.e.*, individual plants); statistical significance of differences was assessed using ordinary one-way ANOVA with p < 0.001, indicated by lowercase letters.

Co-incident with the increasing leaf temperatures, a decline in photosynthetic performance was seen, including depressed Phi2 and F ^’^/F ^’^ values indicative of photoinhibition at PSII (Maxwell and Johnson, 2000; Baker, 2008), and elevated NPQ_t_ values (Tietz et al., 2017) (**Figure 1D, E and F, Supp Table S2**). Furthermore, immunoblotting against Heat Shock Protein of 70 kD (HSP70), a marker of heat stress in plants, demonstrated a marked increase in abundance over the course of the heat stress and a return to pre-stress levels within 3 days of heat recovery (dHR) (**Supp Fig S1D**). Thus, we concluded that our treatment applied an effective heat stress and recovery treatment that is suitable for us to test the role of *Z. mays* plastoglobules in adaptation to heat stress and subsequent recovery.

We selected six time points across the stress and recovery phases of the time-course for detailed investigation of cellular ultrastructure and protein and lipid compositions of isolated plastoglobules and thylakoids: pre-stress (the day prior to the first step-up in temperature; 25 °C), after 1, 3, and 7 days of 40 °C heat stress (dHS), and after 1 and 3 days of heat recovery (dHR) (**Figure 1A**).

### Heat stress effects on *Z. mays* chloroplast ultrastructure

Swelling and/or proliferation of plastoglobules are characteristic responses to both biotic and abiotic stresses across many plant species (van Wijk and Kessler, 2017; Zechmann, 2019; Lundquist et al., 2020). To assess the impact of our heat stress treatment on plastoglobule morphology, and cellular ultrastructure more broadly, we collected transmission electron micrographs (TEMs) of chemically fixed leaf tissue at each of the six time points (**Figure 2, Supp Table S3**).

**Figure 2.**
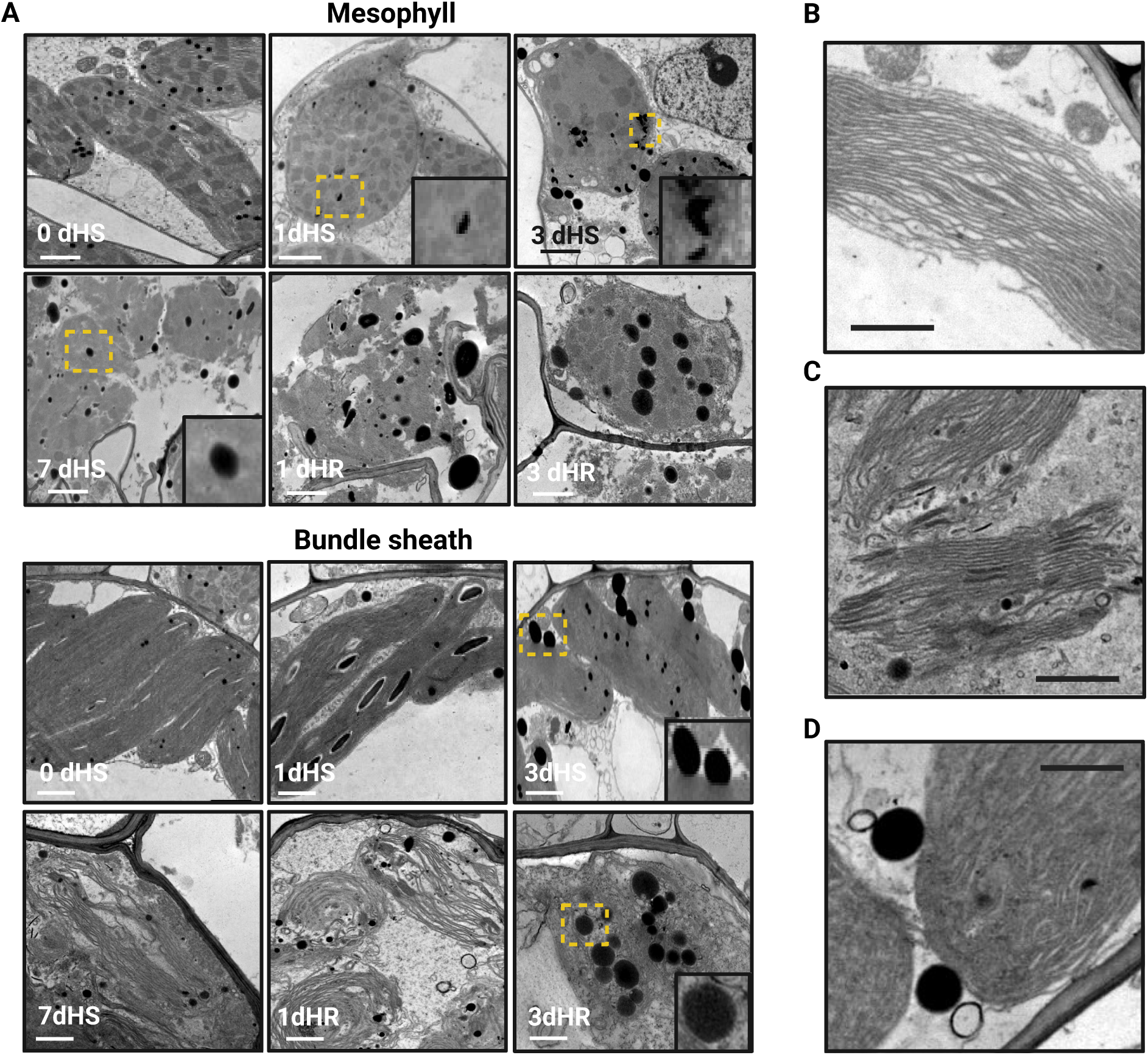
Cellular ultrastructure of mesophyll and bundle sheath cells during the heat stress-and-recovery time-course. **A)** Representative transmission electron micrographs of mesophyll cells (top) and bundle sheath cells (bottom) at each of the six time points of the study. Yellow rectangles demarcate the zoomed regions shown in the inset. **B**) Micrograph of a mesophyll cell at 3 dHS illustrating the ‘swollen thylakoid’ phenotype. **C)** Micrograph of a bundle sheath cell at 7 dHS illustrating the ‘disintegrating chloroplast’ phenotype. **D)** Micrograph of a mesophyll cell at 3 dHS illustrating the presence of cytosolic lipid droplets in close apposition to a chloroplast. Scale bars in all panels designate 1 μm.

Average chloroplast cross-sectional area in mesophyll (M) and bundle sheath (BS) cells declined by about a third over the course of the stress and only partially recovered within 3 days of the return to permissive growth temperatures (**Table 1**). Average plastoglobule size increased in both cell types within one day at 40 °C, and continued to steadily increase throughout the time course, even during the recovery phase, with maximal plastoglobule size appearing at the last time point, 3 dHR (**Table 1**). Likewise, the abundance of plastoglobules (number per µm^2^ of chloroplast cross-sectional area) also increased during the stress in both cell types and continued to increase during the recovery phase (**Figure 2A**). The proliferation of plastoglobules (*i.e.,* increase in the combination of both size and abundance), combined with the decrease in chloroplast size, led to a substantially larger proportion of chloroplast cross-sectional area being occupied by plastoglobule material (plastoglobule area per chloroplast area). Pre-stress, the ratio was distinctly lower in BS cells than M cells, however in both cell-types the ratio increased substantially. It is notable that the proliferation of plastoglobules continued their upward trajectory over the full course of the treatment.

**Table 1.** Ultrastructural measurements of Z. mays chloroplasts and plastoglobules during a heat stress time-coursea.

| Cell Type | Time Point | Chloroplast X-Sectional Area ($\mu\text{m}^2$) | Average Plastoglobule Diameter (nm) | # Plastoglobules / $\mu\text{m}^2$ Chloroplast | Plastoglobule Area / Chloroplast Area (x 1000) |
| --- | --- | --- | --- | --- | --- |
| Bundle Sheath | 0 dHS | 12.8 ± 3.5 | 104.0 ± 7.3 | 0.20 ± 0.06 | 1.7 ± 0.6 |
| | 1 dHS | 14.6 ± 4.2 | 125.0 ± 28.6 | 0.17 ± 0.07 | 2.2 ± 1.5 |
| | 3 dHS | 11.8 ± 2.3 | 136.1 ± 28.3 | 0.33 ± 0.11 | 4.9 ± 2.6 |
| | 7 dHS<sup>b</sup> | 8.3 ± 2.7<sup>b</sup> | 147.5 ± 53.4<sup>b</sup> | 0.39 ± 0.24<sup>b</sup> | 9.2 ± 11.2<sup>b</sup> |
| | 1 dHR | 6.0 ± 0.7 | 138.9 ± 50.7 | 0.58 ± 0.13 | 10.1 ± 8.6 |
| | 3 dHR | 9.4 ± 3.5 | 210.7 ± 38.3 | 0.86 ± 1.24 | 42.4 ± 70.9 |
| Mesophyll | 0 dHS | 15.9 ± 3.6 | 117.7 ± 9.7 | 0.37 ± 0.21 | 4.2 ± 2.6 |
| | 1 dHS | 15.5 ± 5.4 | 134.6 ± 26.3 | 0.29 ± 0.16 | 4.6 ± 3.1 |
| | 3 dHS | 12.8 ± 1.7 | 143.7 ± 14.9 | 0.32 ± 0.15 | 5.4 ± 3.0 |
| | 7 dHS<sup>b</sup> | 10.5 ± 1.5<sup>b</sup> | 175.1 ± 53.9<sup>b</sup> | 0.56 ± 0.26<sup>b</sup> | 12.3 ± 4.2<sup>b</sup> |
| | 1 dHR<sup>b</sup> | 4.5 ± 0.9<sup>b</sup> | 153.1 ± 44.3<sup>b</sup> | 0.46 ± 0.64<sup>b</sup> | 6.3 ± 7.0<sup>b</sup> |
| | 3 dHR | 11.0 ± 3.1 | 195.3 ± 55.1 | 0.77 ± 1.15 | 40.9 ± 73.2 |

<sup>a</sup> Mean $\pm 1$ standard deviation; $n = 4$ biological replicates (four individual plants) with each replicate comprising the average value of a minimum of ten chloroplasts. Only intact chloroplasts fully enclosed within a micrograph were used for measurements.
<sup>b</sup> The prevalence of broken chloroplasts at these time points limited the number of possible measurements. Hence, these values are derived from only 4-7 chloroplasts per replicate and, in the case of the mesophyll cells, only three biological replicates.

We also observed a “crescent” staining pattern of plastoglobules of bundle sheath cells during the stress. While no such crescent-stained plastoglobules were observed at 0 dHS, they were occasionally seen at 1 dHS and became progressively more common over the course of the stress treatment. We have previously reported this staining phenomenon in field-grown *Z. mays* specifically at the R2 developmental stage when plant leaves are transitioning into senescence (Ying et al., 2023). We speculated that this staining pattern may reflect partial turnover of the plastoglobules, with turnover initiating on one side and progressing in a directional manner. If correct, this would suggest an increasing rate of plastoglobule turnover during our heat stress treatment, that has initiated within the first day of heat stress.

In addition to the effects on plastoglobule morphology, clear effects could be seen among the thylakoids, which demonstrated a progressively greater frequency of swelling between thylakoid membranes of the granal stacks in M cells (**Figure 2B**). At peak stress (7 dHS), though not at the earlier stress timepoints, chloroplasts typically appeared to have lost their integrity, with envelopes largely or fully disintegrated and thylakoid membranes and other chloroplast material ‘spilled’ into the cytosol (**Figure 2C**). This sensitivity of chloroplast integrity appeared to be especially acute among M cells, where intact chloroplasts were especially rare. Within 1 day after recovery (1 dHR) chloroplasts of both cell types generally appeared disrupted, as seen at 7 dHS, although by 3 dHR chloroplast integrity had largely recovered. Finally, consistent with observations of other groups (Mueller et al., 2015; Mueller et al., 2017; Scholz et al., 2025), we observed increasingly frequent cytosolic dark-staining lipid droplets in both M and BS during the heat stress, characterized by larger diameters (*ca.* 400 – 800 nm) than the plastoglobules, although their strong osmiophilicity was indistinguishable from that of the plastoglobules. The cytosolic lipid droplets regularly appeared to be in physical contact with chloroplast envelope membranes (**Figure 2D**). Remarkably, cytosolic lipid droplets continued to accumulate within 1 dHR and were often seen in even larger sizes of 1 -2 µm in diameter. We also occasionally noted the crescent-staining pattern of cytosolic lipid droplets, similar to what was seen among some plastoglobules, which were always adjacent to chloroplasts (**Supp Fig S2**). Collectively, our ultrastructural results document maize leaf tissue undergoing progressively more severe effects on the chloroplast ultrastructure, including a clear proliferation of plastoglobules, contemporaneously with the emergence of prevalent cytosolic lipid droplets.

### Total leaf proteomics

To assess the impact of the heat stress treatment on leaf tissue we carried out a proteomic investigation of whole leaf tissue at the pre-stress (0 dHS), peak stress (7 dHS), and recovery (3 dHR) time-points. Proteins were in-gel digested and analyzed using nano-liquid chromatography connected in-line with a Q-Exactive HF tandem mass spectrometer (nLC-MS/MS). Proteins were quantified using label-free quantification, as implemented in the MaxQuant software program (Cox et al., 2014), and normalized to total LFQ intensity (nLFQ); in this way, the nLFQ intensity corresponds to the relative proportion of protein mass within each sample. At a 1% estimated false discovery rate at the peptide and protein levels, we identified 1049 proteins or protein groups (hereafter, proteins) across the three time points (**Supp Table S4**). As expected, many heat shock proteins and related protein chaperones across many sub-cellular compartments increased substantially from 0 to 7 dHS and were partially recovered upon removal from the heat stress (**Supp Fig S3A, Supp Table S4**).

We assessed the protein contribution of sub-cellular compartments in the leaf samples. Consistent with the shrinking chloroplast size observed by TEM, the proportion of chloroplast proteins was reduced *ca.* 17% from 0 to 7 dHS, while the vacuole, cytosol, and mitochondria were significantly increased (**Figure 3A, Supp Fig S3B**). A partial recovery of heat shock protein levels was seen by 3 dHR.

**Figure 3.**
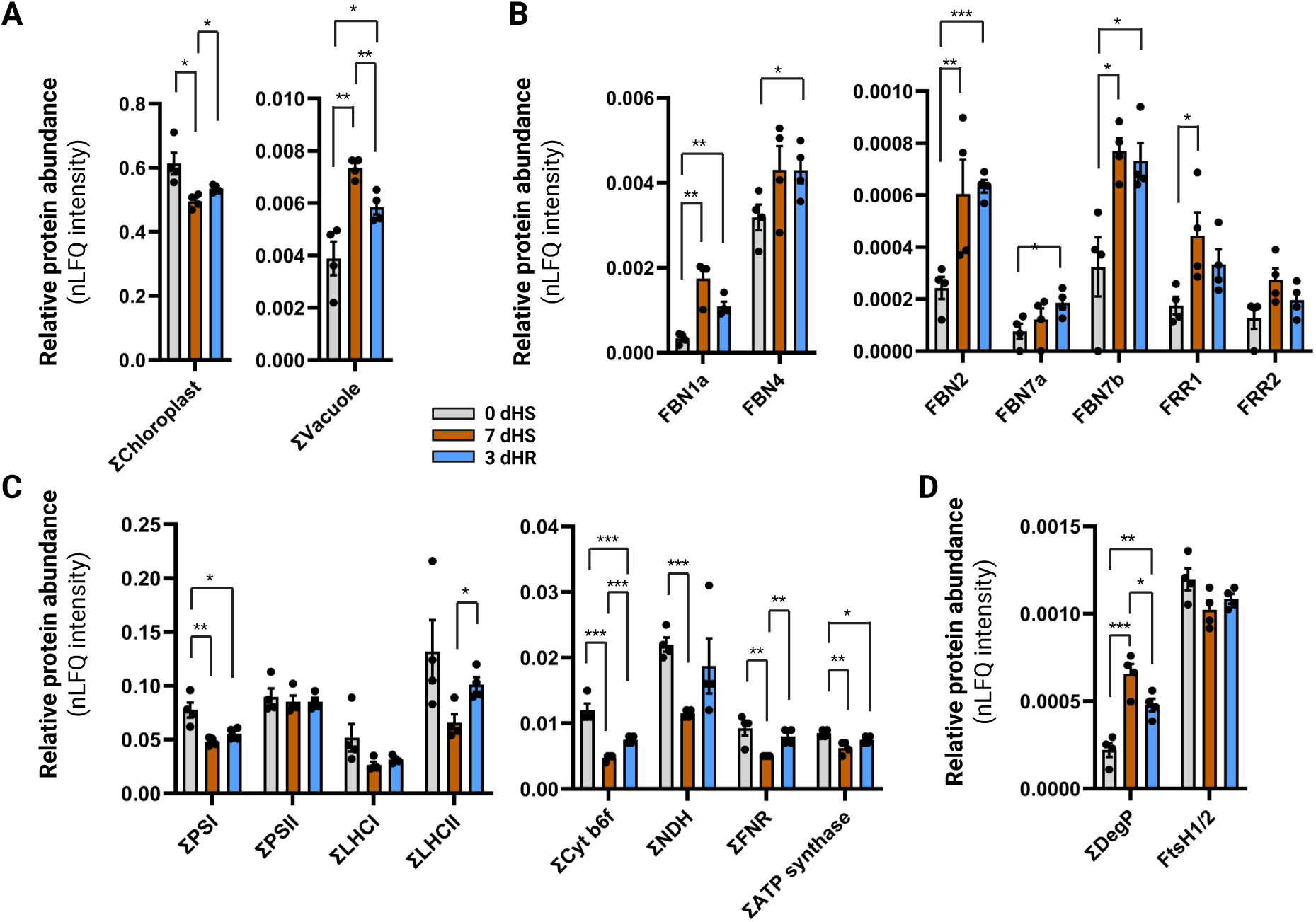
Quantitative proteomics of *Z. mays* leaf tissue at three time points during the heat-stress-and-recovery time-course. **A**, Protein abundance of selected sub-cellular compartments from total leaf protein samples. nLFQ intensities of all proteins annotated with a localization in the respective sub-compartment, as annotated in the Plant Proteome Database, were summed. **B**, Relative protein abundance of selected plastoglobule proteins. **C**, Relative protein abundance of selected thylakoid-localized photosynthetic complexes. nLFQ intensities of all proteins annotated as a part of the respective complex according to the Plant Proteome Database were summed. **D**, Relative protein abundance of selected chloroplast proteases. nLFQ intensities of all proteins annotated as a part of the respective complex according to the Plant Proteome Database were summed. Data are represented as bar graphs plotting the mean ± 1 s.e.m, n = 4 biological replicates. Individual data points are indicated as black dots. * p < 0.05, ** p < 0.01, *** p < 0.001, homoscedastic two-tailed Student’s t-test.

Consistent with the proliferation of plastoglobules observed by TEM, the plastoglobule protein, Fibrillin 1a (FBN1a), increased over 5-fold between 0 and 7 dHS (**Figure 3B**). Several other plastoglobule proteins [FBN2, FBN7a, Flavin reductase-related 1 (FRR1) and FRR2] also increased by 2- to 3-fold under the stress, although these changes occurred with less statistical significance, likely due to their low abundance in the total leaf samples (**Figure 3B**). In contrast to the heat shock proteins, these plastoglobule proteins tended to remain at elevated abundance during the recovery phase, consistent with the persistently high levels of plastoglobules.

Photosynthetic complexes of the light-harvesting reactions broadly declined under the stress and partially recovered after removing the stress (**Figure 3C**). This decrease was especially conspicuous for the Cytochrome b_6_f complex, the NAD(P)H dehydrogenase complex, and the Ferredoxin-NAD(P)H reductase complex. Photosystem II was a curious exception, since its protein levels remained constant over the time points (**Figure 3C**), although the D1 protein (PsbA) levels declined by 2-fold (**Supp Table S4**), aligning with the substantial drop in F_v_’/F_m_’. The FtsH1/2/5/8 complex has been associated with the turnover of the D1 protein, as well as other proteins of photosynthesis (Yu et al., 2005; Zaltsman et al., 2005; van Wijk, 2024). However, levels of the FtsH1 and FtsH2 did not increase under the stress while FtsH5 and FtsH8 were not identified in our proteomic analysis, presumably present but below the detection limit of our experimental system (**Figure 3D**). In contrast, the proteins of the DegP protease complex (Sun et al., 2007; van Wijk, 2015) were significantly increased during the stress and only partially recovered by 3 dHR (**Figure 3D**).

### Isolation of plastoglobules and thylakoids from maize leaf tissue

Thylakoids and plastoglobules were isolated from leaf tissue at each of the six time points of the heat treatment to allow a close, parallel investigation of the composition of thylakoids and plastoglobules. Accordingly, thylakoids and plastoglobules were isolated from the same leaf tissue at each of the six time points of the heat treatment and subsequent recoveries. Consistent with the observed increase in plastoglobule size and number seen in the TEMs, the yield of isolated plastoglobule material increased significantly over the time-course of heat stress and recovery (**Figure 4A & 4B**). To avoid artefactual effects on the thylakoids, we did not sonicate the isolated thylakoids, resulting in the co-isolation of physically associated plastoglobules. This was considered to have minimal impact on our study due to the overwhelming quantity of thylakoid material relative to that of the associated plastoglobules. Immunoblotting for marker proteins (PsbA [D1] for thylakoids and Fibrillin1a [FBN1a] for plastoglobules) confirmed effective enrichment of both sub-compartments (**Figure 4C**). Consistent with the proliferation of plastoglobules, FBN1a protein levels in the isolated thylakoids increased progressively over the seven days of the heat stress (**Figure 4C**).

**Figure 4.**
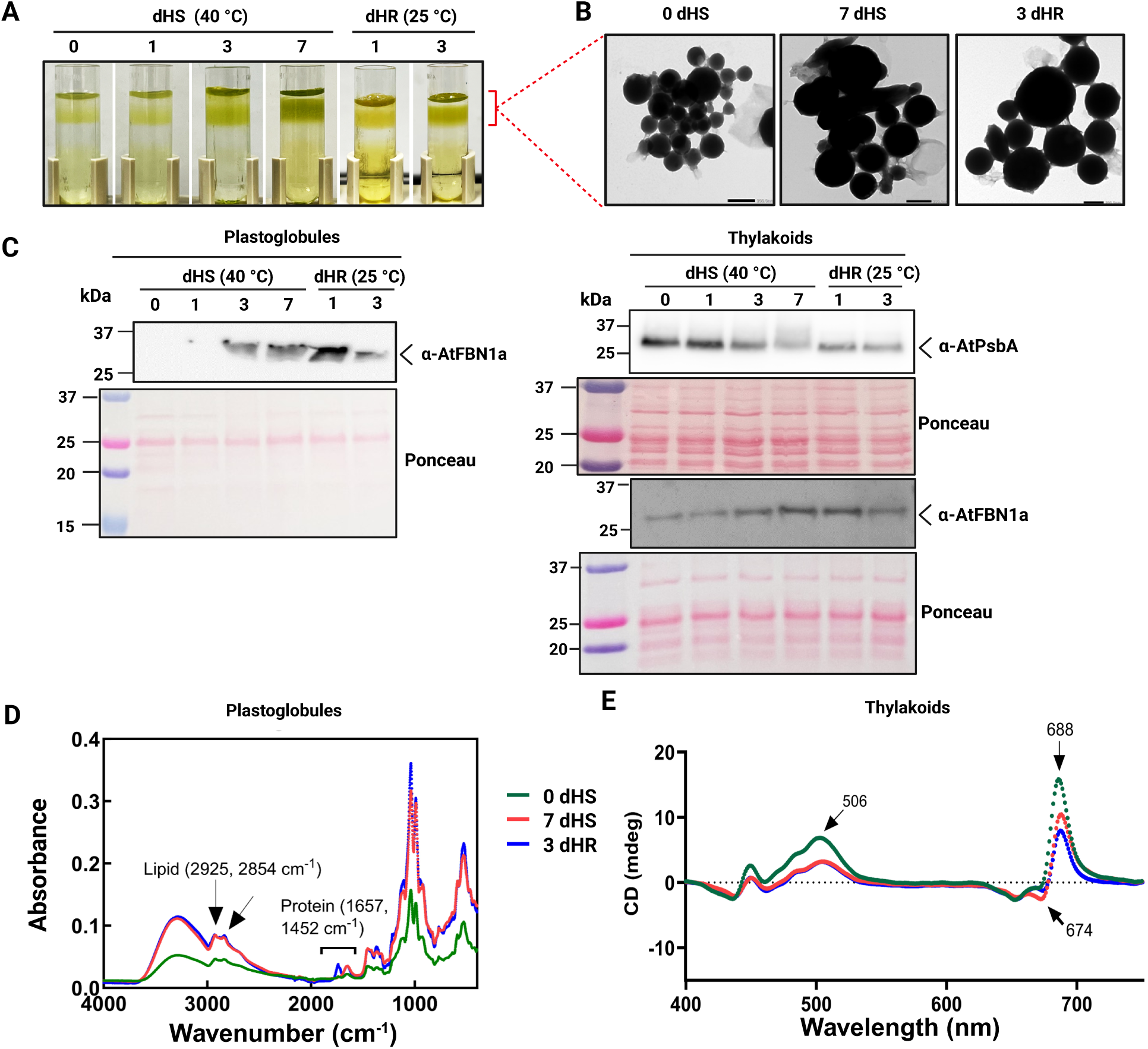
Isolation of *Z. mays* plastoglobules and thylakoids during the heat-stress-and-recovery time-course. **A)** Plastoglobule isolation following flotation on a sucrose gradient. Plastoglobules are found as a yellow pad at or near the surface of the gradient, indicated with the red bracket. **B)** Transmission electron micrographs of purified plastoglobules, extracted from the top of the sucrose gradient, at the three specified time points. Black scale bars = 200 nm. **C)** Immunoblots of isolated plastoglobules (left) and thylakoids (right) probed with anti-AtPsbA (D1 protein of PSII core) (anti-Rubisco small subunit as a loading control) and anti-AtFBN1a (Fibrillin 1a) antibodies (marker of thylakoid and plastoglobule, respectively) were used to demonstrate the level of purity of isolations at each time point. **D)** Fourier Transform Infrared (FTIR) spectroscopy was used to analyze lipid and protein accumulation in isolated plastoglobules at the three designated time points. Prior to the spectroscopic measurements, plastoglobule samples were normalized to an OD_700_ of 1.0. **E)** Disorganization of isolated thylakoids was assessed using Circular Dichroism (CD) at 688 nm, showing a major peak at PSII.

Isolated plastoglobules were analyzed using Fourier Transform Infrared (FT-IR) spectroscopy. Notable lipid signals, identified by peaks at 2925 cm□¹ and 2854 cm□¹, along with protein content signals at 1652 cm□¹ and 1452 cm□¹ (Lahlali et al., 2014), were substantially more pronounced under both heat stress (7 dHS) and recovery (3 dHR) conditions compared to pre-stress (0 dHS) conditions (**Figure 4D**). This is consistent with the presence of a specific population of proteins associated with the *Z. mays* plastoglobules, corroborated by the proteomic analyses described below. Moreover, the elevated ratios of protein-associated bands to lipid-associated bands at 0 dHS indicate a higher protein:lipid ratio pre-stress, consistent with their smaller diameters that would accommodate more surface-associated protein per unit of lipid.

Circular dichroism (CD) was used to characterize the organization of the thylakoid membrane pigments. A decline was observed in the positive band at 688 nm and the negative band around 674 nm, both of which are linked to Chl *a* molecules associated with the chiral macrodomains of photosynthetic complexes (Toth et al., 2016). Additionally, the band near 506 nm, attributed to carotenoids in a long-range ordered structure, showed significant differences between pre-stress (0 dHS) conditions and those exposed to heat stress (7 dHS) or heat recovery (3 dHR) in maize leaf thylakoids (**Figure 4E**). These observations are indicative of a reduction of photosystem structure within the isolated thylakoids from heat-stressed maize which persists at least until 3 dHR, consistent with our TEM chloroplast ultrastructure data (**Figure 2**). Subsequently, isolated thylakoids and plastoglobules were subjected to detailed proteomic and lipidomic characterization, as described below.

### Effects on the Thylakoid and Plastoglobule Proteomes

Quantitative proteomics was performed with the isolated thylakoids and plastoglobules at all six time points of the heat stress treatment using the same methodology as for the total leaf samples. Across all time points, a total of 734 proteins were identified in the thylakoid samples and 1261 in the plastoglobule samples (**Supp Table S5 & S6**).

The quantitative thylakoid proteome was remarkably stable during the time course (**Figure 5A**). Statistically significant changes in abundance were limited to only about 40 proteins. Notably, effects were most prominent at the 3 dHS time point, not the more extreme physiological state experienced at 7 dHS. In view of the ultrastructure results, which demonstrated pervasive disassembly of thylakoids by 7 dHS, we suspect that the 3 dHS time point represents thylakoids that are encountering the heat stress but remain intact and recoverable, while at 7 dHS much of the thylakoid material has progressed to a sufficiently disassembled state that they are not recoverable by our isolation method and we predominantly isolate those thylakoids that have ‘escaped’ the stress and remain intact. At 1 dHS, the thylakoid proteome was generally unperturbed; only a single protein, the Cytochrome f subunit of the Cytochrome b_6_f complex was decreased with statistical significance at this time point (by *ca.* 50% relative to 0 dHS) (**Supp Table S5**). By 3 dHS this subunit has decreased further, and 29 other proteins are also changed with statistical significance. Not surprisingly, various subunits of each of the photosynthetic complexes were substantially reduced at this time point. A number of proteins were also increased in abundance including the PsbS subunit that mediates the energy-dependent form of NPQ (Li et al., 2000), 13-Lipoxygenase 10 and/or 11 (LOX10/11), Allene oxide synthase 2b, and multiple proteins of the Calvin-Benson cycle that are ostensibly localized in the stroma, including RubisCO large and small subunits (RbcL and RbcS, respectively), Phopshoglycerate kinase (PGK), Glyceraldehyde-3-phosphate dehydrogenase (GAPDH), and Transketolase (TKL) (**Figure 5B, Supp Fig S4, Supp Table S5**). Notably, the Calvin-Benson cycle proteins, as well as LOX10/11 and AOS2b, are not increased in abundance at the total leaf level (in fact, RbcL is significantly decreased), despite substantially increased abundance of each of these proteins at the thylakoids and plastoglobules, although our total leaf analyses does not include the 3 dHS time point when abundance at the plastoglobule and thylakoid are typically highest. This increased abundance at thylakoids and plastoglobules is uniformly reversed within 1 dHR. This suggests that they are re-mobilized to the thylakoids and/or plastoglobules under the heat stress and rapidly released from these sub-compartments upon removal of the heat stress. Localization of multiple Calvin-Benson cycle enzymes at plastoglobules would align with the validated plastoglobule localization of Fructose Bis-phosphate Aldolase to *A. thaliana* plastoglobules (Vidi et al., 2006; Ytterberg et al., 2006). From our results it cannot be discerned whether the increased abundance of these enzymes reflects their localization at the thylakoids, plastoglobules, or both. However, the fact that the changes in abundance under stress are mirrored between the thylakoids (which retain attached plastoglobules) and the plastoglobules, suggests that they are predominantly remobilizing to the plastoglobules. Furthermore, there is precedent for the remobilization of a biosynthetic pathway (*i.e.*, the jasmonic acid pathway) to plastoglobules under stress in *A. thaliana* (Lundquist et al., 2013). Notably, although increased levels of the Calvin-Benson cycle enzymes were largely or fully reversed within 1 dHR, the decreased levels of the photosynthetic subunits were only partially reversed by 3 dHR. This indicates a more rapid restoration of the Calvin-Benson cycle enzymes than the photosynthetic machinery under the recovery phase of the time-course.

**Figure 5.**
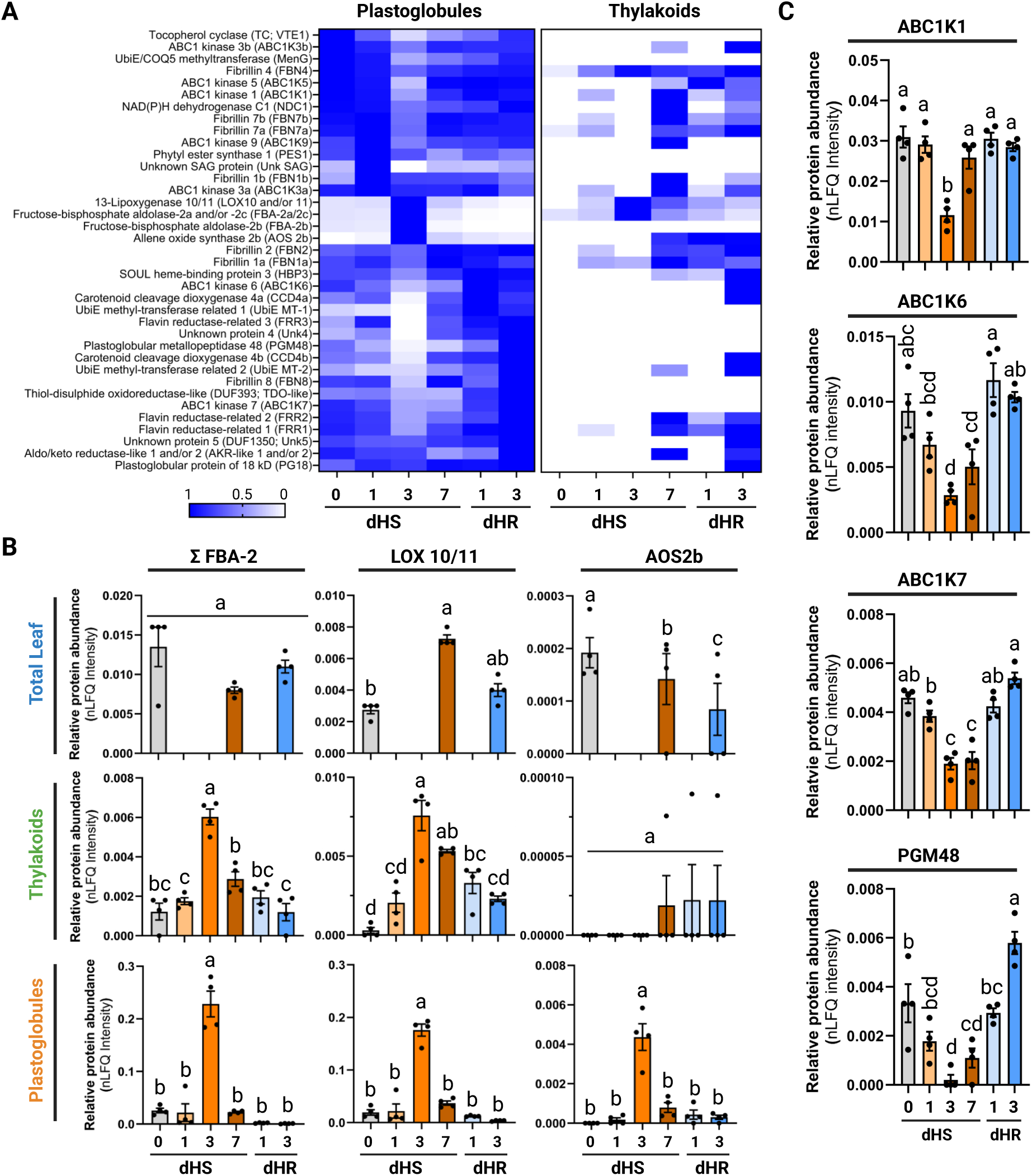
Quantitative proteomics of isolated *Z. mays* plastoglobules and thylakoids during the heat-stress-and-recovery time-course. **A)** Heatmap of normalized nLFQ intensities of all 34 ‘core’ proteins of the *Z. mays* plastoglobule proteome. Each protein is normalized to its maximal protein abundance within each sample, such that each protein will have its peak abundance of 1.0 at one of the six time points. Proteins are sorted by the time point at which they hold maximal protein abundance in the plastoglobule samples. **B)** Relative protein abundance of the summed levels of Fructose-bisphosphate aldolase 2 isoforms, 13-Lipoxygenase 10 and/or 11 (LOX 10/11), and Allene oxide synthase 2b (AOS2b), in total leaf, thylakoids and plastoglobules. Note that the total leaf proteome was assessed at only three of the time points. **C)** Relative protein abundance of ABC1K1, ABC1K6, ABC1K7, and PGM48 in plastoglobules. Statistical significance of differences in panels B and C were assessed using ordinary one-way ANOVA with p < 0.001, indicated by lowercase letters. Graphs plot the mean ± 1 s.e.m., n = 4 biological replicates.

The use of highly sensitive LC-MS/MS permits identification of even very low abundant contaminant proteins in our samples; indeed, the lowest abundant 793 proteins in our plastoglobule samples cumulatively amounted to only 1% of the total nLFQ intensity. Thus, to distinguish *bona fide* plastoglobule proteins from contaminants, we relied on homology to the plastoglobule proteome of *A. thaliana*, which has been carefully curated via multiple, comparative proteomic studies, fluorescent protein tagging and immunogold labelling across multiple research groups (Vidi et al., 2006; Ytterberg et al., 2006; Lundquist et al., 2012; Espinoza-Corral et al., 2021). This is the same approach that we have used in our associated companion paper investigating the *Z. mays* plastoglobules under a water-deficit stress (Devadasu, et al. unpublished). Clear homologs of each of the *A. thaliana* plastoglobule proteins are encoded in the maize genome, all but four of which were identified in our plastoglobule isolations (**Table 2, Supp Table S7**). The exceptions were the stress-induced Esterase/Lipase/Thioesterase 4 (ZmELT4, GRMZM5G817559), also missing from the water-deficit dataset, and one plastoglobule protein (ZmNDC1b) that was identified in the water-deficit plastoglobule proteome (Devadasu, et al. unpublished), but not the heat stress plastoglobule proteome described here. Thus, we defined a set of 37 proteins comprising the *Z. mays* plastoglobule proteome from our heat stress experiment (**Table 2, Supp Table S7**).

**Table 2.**
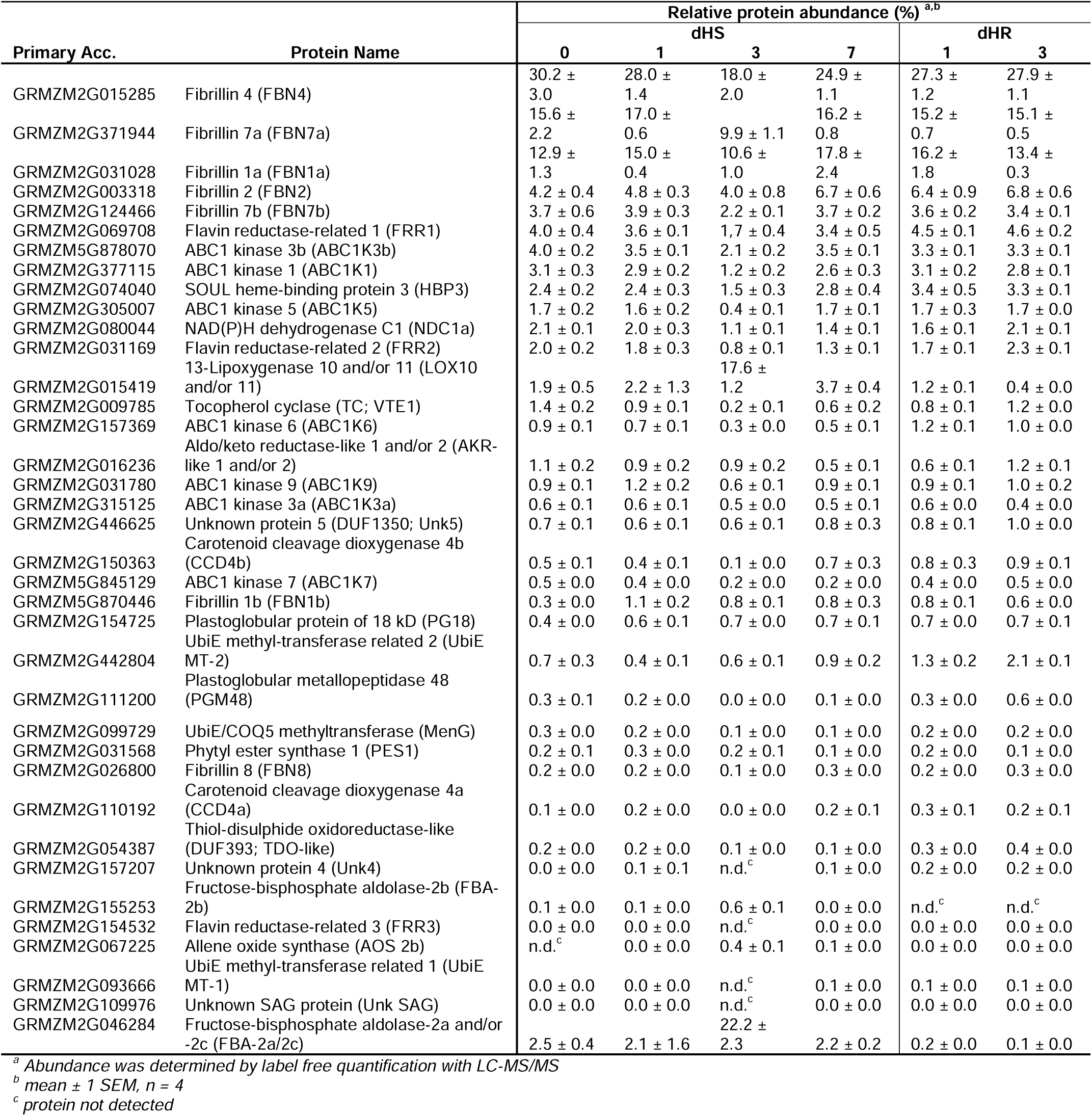
The quantitative Z. mays plastoglobule proteome under heat stress.

The quantitative composition of the plastoglobule proteome was broadly similar to that seen under water-deficit stress, however the proteome was impacted more extensively by the heat stress (**Figure 5A**). As for the thylakoids, the effects were most prominent at the 3 dHS time point, not 7 dHS. Nearly half of the proteome, 15 proteins, were changed with statistical significance at 3 dHS relative to 0 dHS, all of which were reversed within 1 dHR (**Table 2, Supp Table S7**). Among these proteins were five of the seven plastoglobular ABC1K orthologs, which were reduced between 2- and 4-fold. Notably the reduction of ABC1K6 and ABC1K7 persisted into 7 dHS, whereas the other ABC1Ks returned to pre-stress levels at 7 dHS (**Figure 5C**), consistent with the parallel accumulation patterns of ABC1K6 and ABC1K7 observed previously in *A. thaliana* (Espinoza-Corral and Lundquist, 2022). Also found in this group were both Flavin reductase-related isoforms (FRR1 and 2), which were also reduced over 2-fold, Plastoglobular metallopeptidase M48 (PGM48; reduced over 16-fold) and Tocopherol cyclase (TC; reduced over 4-fold) (**Figure 5C**, **Table 2, Supp Table S7**). These were countered by increases in the levels of LOX10/11, Allene oxide synthase 2b (AOS 2b), and the FBA2 isoforms, as noted above (**Figure 5B**). Note that although evidence of remobilization of other Calvin-Benson cycle enzymes, beyond the FBA 2 isoforms, was observed in our thylakoid and plastoglobule proteome analysis, we do not include these proteins in our ‘core’ plastoglobule proteome represented in **Table 2**, because they have not previously been identified as plastoglobule-localized with more rigorous determination.

Notably, all plastoglobular FBNs remained relatively stable over the heat stress time-course. This contrasts with the water-deficit stress in which FBN4 was significantly reduced during the stress and FBN1a and FBN1b were significantly increased. It is also notable that the Unknown protein senescence-associated gene (UnkSAG), which was increased substantially under water-deficit stress, showed only a muted increase during our heat stress, and limited to the 1 dHS time point. In fact, statistically significant changes to the plastoglobule proteome that were shared across both stress treatments were limited to the increased levels of LOX10/11 and FBA2 isoforms. Pointedly, the apparent remobilization of additional Calvin-Benson cycle enzymes was not reflected under the water-deficit stress. This suggests that different role(s) are employed by *Z. mays* plastoglobules when faced with a water-deficit *versus* a heat stress treatment.

### Thylakoid Lipid Remodeling in Response to Heat Stress

Membrane remodeling is a prominent component of plant responses to temperature fluctuations. To understand the nature and extent of membrane remodeling in the *Z. mays* thylakoids over the course of our heat stress treatment we employed thin layer chromatography and gas chromatography with flame ionization detection (GC-FID) to quantitatively profile polar lipids (**Supp Tables S8 & S9**). Consistent with the observed disassembly of thylakoids observed by TEM, total thylakoid lipids decreased between 1 and 3 dHS and continued to decline between 3 and 7 dHS, reaching less than half of the pre-stressed levels (**Figure 6A**). These levels did not recover to an appreciable amount during the 3 days of HR. Thylakoid lipid material was dominated by C18:3 acyl chains but decreased from approximately 83 mol% of total thylakoid lipid at 0 dHS, to approximately 62 mol% at 7 dHS. The reduction in C18:3 was offset by increases in C16:0, C18:0 and C18:2 (**Figure 6B**). Thylakoids were comprised of MGDG, DGDG, SQDG and PG. Only low levels of contaminating lipids, including phosphatidylcholine, phosphatidylethanolamine, and phosphatidylinositol were found in our isolated thylakoids (**Supp Table S9**). As expected, the galactolipids accounted for the vast majority of polar lipids, however, the levels of MGDG decreased substantially under stress, whereas levels of DGDG increased. This manifested as a sharp drop in the MGDG:DGDG ratio, from over 3-to-1 pre-stress, to about 0.5-to-1 at peak stress, which only partially recovered by 3 dHR (**Figure 6C & D**).

**Figure 6.**
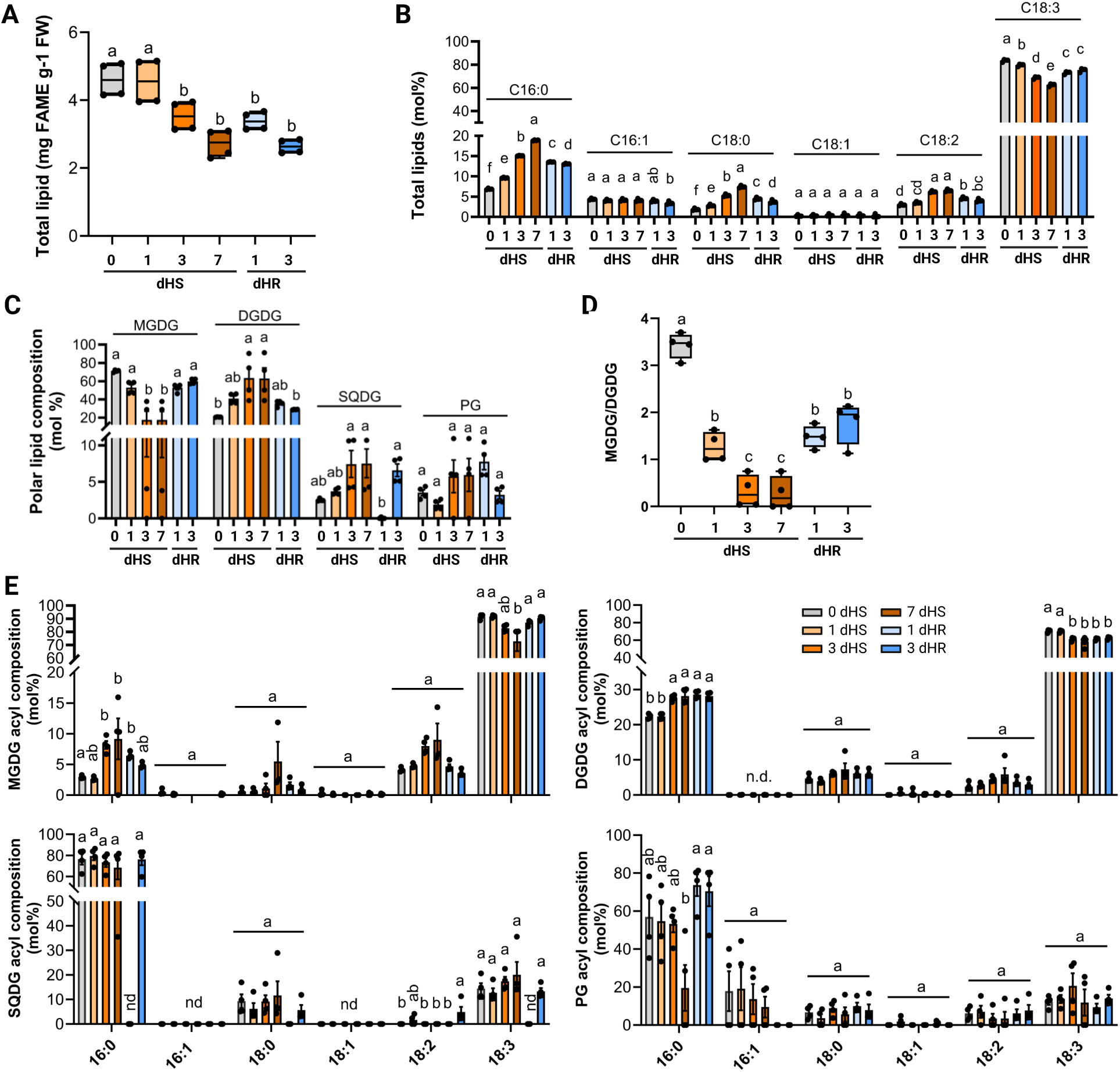
Lipidome profiling of isolated *Z. mays* thylakoids during the heat-stress-and-recovery time-course. **A)** Total lipid content from isolated thylakoids, measured as fatty acid methyl esters (FAMEs) derived from saponification and methylation of acyl groups. **B)** Total lipid acyl chain composition. Note that C16:1 includes both cis and trans isomers and that C16:2 and C16:3 acyl chains were not detected in any samples and are not plotted in this chart. **C**) Polar lipid compositions of each lipid class. Lipids were separated by thin layer chromatography, scraped from the plates, saponified and analyzed by gas-liquid chromatography with flame ionization detection. Note that C16:2 and C16:3 acyl chains were not detected in any samples and are not plotted in this chart. **D**) Ratio of MGDG to DGDG at each time point measured on a mole basis. **E**) Acyl group distributions of monogalactosyl diacylglycerol (MGDG), digalactosyl diacylglycerol (DGDG), sulfoquinovosyl diacylglycerol (SQDG), phosphatidylglycerol (PG). Note that SQDG was not detected at the 1 dHR time point. Relative molar ratios of each polar lipid class are plotted as bar plots with individual data points overlaid as black dots. Statistically significant groupings between time points (and within a polar lipid class) are indicated in all panels by letter codes using ordinary two-way ANOVA, p < 0.001, n = 4 biological replicates.

The acyl composition of MGDG shifted substantially towards higher saturation under the stress, with a decrease of C18:3 from about 90 mol% to 70 mol%, which was offset by increases in C16:0, C18:0 and C18:2 (**Figure 6E**). A similar trend was seen with the DGDG lipids, albeit more muted. Notably, these shifts in the acyl composition in MGDG and DGDG were manifested primarily between the 1 and 3 dHS timepoints and only partially (MGDG), or did not (DGDG), recover within 3 dHR (**Figure 6E**). In contrast, acyl composition of SQDG, which was predominantly comprising C16:0, was unaffected by the stress and PG shifted slightly (albeit without statistical significance) towards less saturated acyl chains. Broadly speaking, we see that the thylakoid membrane responded to the heat stress by reducing the MGDG:DGDG ratio and increasing the level of saturated acyl chains.

### LC-MS/MS-based Lipidomics of Thylakoids and Plastoglobules

We employed LC-MS/MS-based lipidomics to interrogate the thylakoid and plastoglobule lipidomes at each time point. A total of 130 distinct compounds were annotated in thylakoids and 100 in plastoglobules based on accurate mass matches, retention times, authentic standards and comparison with our previous LC-MS/MS lipidome study of *Synechocystis* sp. PCC6803 cyanoglobules (**Supplementary Tables S10 & S11**; Susanto, et al. 2025). As expected, thylakoid lipids were dominated by MGDG and DGDG. Pheophytin showed the single highest MS1 ion intensity among annotated lipids at 0 dHS; coupled with our inability to detect chlorophyll a, we suggest that Mg^2+^ was de-chelated from chlorophyll a during handling, resulting in conversion to pheophytin. This is consistent with the presence of pheophytin, but lack of any detectable chlorophyll a, in the isolated plastoglobules.

In both thylakoids and plastoglobules, triacylglycerol (TAG) levels increased over 10-fold, and diacylglycerol (DAG) levels over 2-fold, during the heat stress and largely returned to pre-stress levels during the 3 days of heat recovery (**Figure 7A, Supp Fig 5A**). We note that low levels of monoacylglycerol were also annotated in thylakoids and plastoglobules, however their levels tended to decline during the stress. Although TAG was clearly observed in our isolated thylakoids, most of these lipids presumably exist in the associated plastoglobules, because they would be only poorly accommodated within the protein-dense thylakoid bilayer. Consistently, these TAG species are highly enriched in the isolated plastoglobules relative to thylakoids (over 19% of total MS1 ion intensity in plastoglobules, versus 6% in thylakoids (**Table 3, Supplementary Table S12**). Prior to stress, TAG species were comprised primarily of 50:0, 48:0, and 52:0, however, this shifted to highly unsaturated acyl groups at peak stress, with the most abundant TAG species being 54:9, 54:8 and 52:6 (**Supplementary Tables S10 & S11**). The preponderance of 54:8 and 54:9 TAG species aligns with the high abundance we observed in *Z. mays* plastoglobules under drought stress in our companion paper (Devadasu, et al. unpublished). At 3 and 7 dHS, the cumulative TAG level in plastoglobule samples had a higher MS1 ion intensity than any other individual or cumulative lipid groups, indicating that TAG is a major component of the plastoglobules, as seen also under drought stress (Devadasu, et al. unpublished). DAG appeared to be much lower abundant in thylakoids and plastoglobules (based on the MS1 ion intensities), although it also increased over 3-fold from 0 to 7 dHS in the plastoglobules. Prior to stress, DAG was predominantly in the 36:0 form but largely shifted to 36:6 and 36:5 during heat stress, consistent with the shift seen with TAG from saturated to polyunsaturated acyl groups. The polyunsaturated acyl composition of the TAG and DAG species under heat stress is consistent with the presence of 18:3 acyl group indicating that the TAG deposited in the plastoglobules under heat stress is derived from the chloroplast membrane lipids.

**Figure 7.**
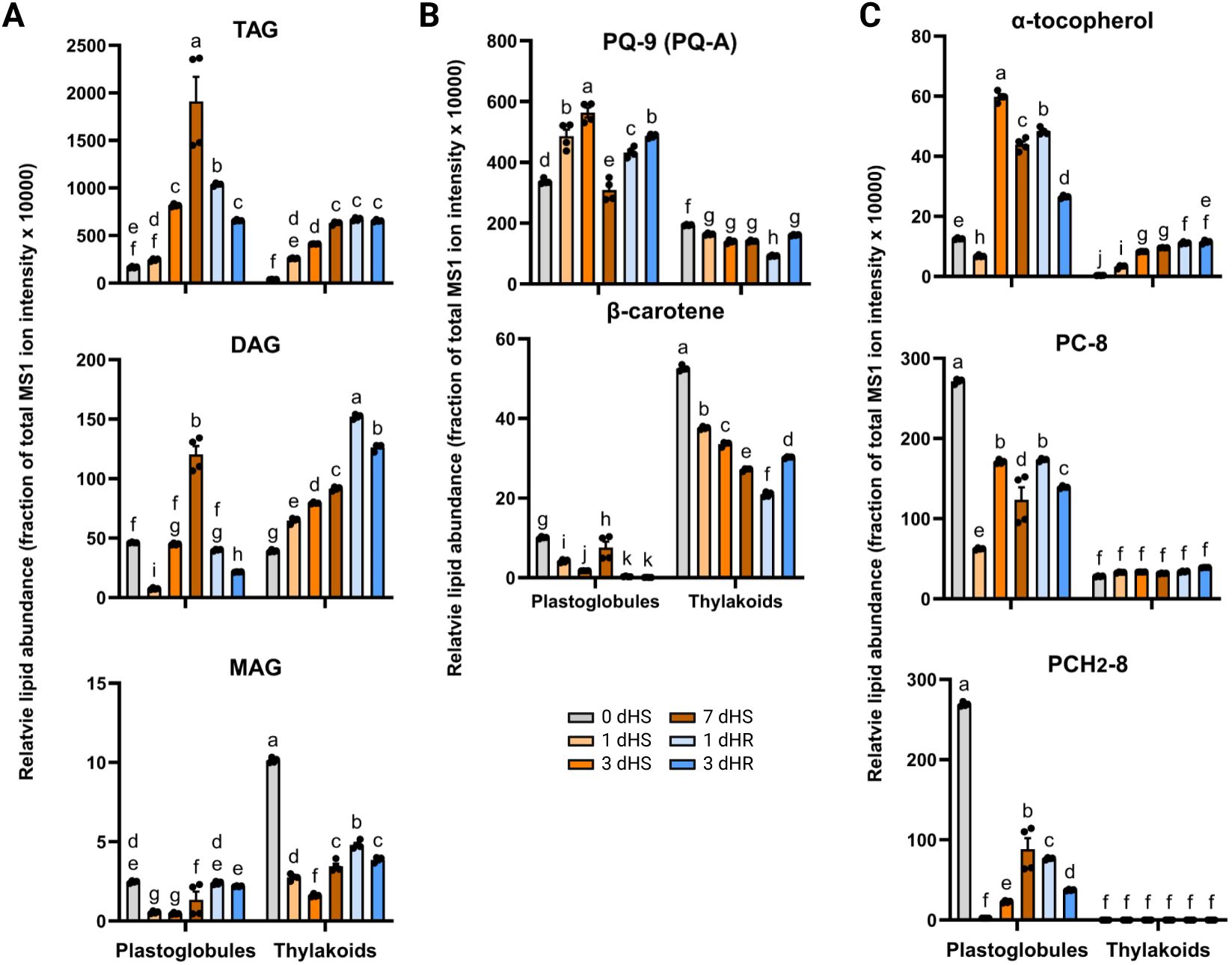
Relative levels of selected lipid compounds from *Z. mays* plastoglobules and thylakoids using LC-MS/MS. Cumulative levels of the neutral lipids use the summed MS1 ion intensities from all acyl chain combinations. Levels are plotted as normalized MS1 ion intensity (x10000) at each of the six time points as a bar chart with individual data points as black dots. Error bars indicate mean ± 1 s.e.m. and are not visible when they are smaller than the individual data points. n = 4 biological replicates. Triacylglycerol, TAG; diacylglycerol, DAG; monoacylglycerol, MAG; plastoquinone-9, PQ-9/PQ-A]); plastochromenol-8, PC-8; plastochromanol-8, PCH_2_-8. Statistical significance of differences was assessed using ordinary two-way ANOVA with p < 0.001, indicated by lowercase letters.

**Table 3.** Relative levels of triacylglycerol species (TAGs) in plastoglobules at each time point of heat stress.

| Acyl composition | 0 dHS | 1 dHS | 3 dHS | 7 dHS | 1 dHR | 3 dHR |
| --- | --- | --- | --- | --- | --- | --- |
| 48:0<sup>a</sup> | 15.7 ± 0.2<sup>b</sup> | 1.3 ± 0.0 | 2.4 ± 0.3 | 2.0 ± 0.5 | 0.8 ± 0.0 | 3.4 ± 0.1 |
| 48:1 | 4.1 ± 0.1 | 0.9 ± 0.0 | 0.2 ± 0.0 | 0.0 ± 0.0 | 0.1 ± 0.0 | 0.2 ± 0.0 |
| 48:2 | 2.4 ± 0.1 | 0.5 ± 0.0 | 0.1 ± 0.0 | 0.0 ± 0.0 | 0.0 ± 0.0 | 0.1 ± 0.0 |
| 48:3 | 0.5 ± 0.0 | 0.2 ± 0.0 | 0.0 ± 0.0 | 0.0 ± 0.0 | 0.0 ± 0.0 | 0.0 ± 0.0 |
| 48:4 | 0.2 ± 0.0 | 0.0 ± 0.0 | 0.0 ± 0.0 | 0.0 ± 0.0 | 0.0 ± 0.0 | 0.0 ± 0.0 |
| 50:0 | 21.7 ± 0.2 | 0.0 ± 0.0 | 4.1 ± 0.6 | 3.2 ± 0.7 | 1.4 ± 0.1 | 5.7 ± 0.2 |
| 50:1 | 3.7 ± 0.2 | 0.7 ± 0.0 | 0.2 ± 0.0 | 0.1 ± 0.0 | 0.1 ± 0.0 | 0.3 ± 0.0 |
| 50:2 | 3.9 ± 0.1 | 1.0 ± 0.0 | 0.4 ± 0.0 | 0.3 ± 0.0 | 0.5 ± 0.0 | 0.6 ± 0.0 |
| 50:3 | 1.4 ± 0.0 | 1.1 ± 0.0 | 1.2 ± 0.0 | 1.3 ± 0.0 | 1.4 ± 0.0 | 1.2 ± 0.0 |
| 50:4 | 0.0 ± 0.0 | 0.7 ± 0.0 | 0.0 ± 0.0 | 0.1 ± 0.0 | 0.1 ± 0.0 | 0.0 ± 0.0 |
| 50:5 | 0.2 ± 0.0 | 0.0 ± 0.0 | 0.0 ± 0.0 | 0.0 ± 0.0 | 0.0 ± 0.0 | 0.0 ± 0.0 |
| 50:6 | 0.1 ± 0.0 | 0.1 ± 0.0 | 0.1 ± 0.0 | 0.2 ± 0.0 | 0.1 ± 0.0 | 0.1 ± 0.0 |
| 52:0 | 13.6 ± 0.2 | 0.3 ± 0.0 | 2.7 ± 0.4 | 2.1 ± 0.5 | 1.0 ± 0.0 | 3.8 ± 0.1 |
| 52:1 | 0.9 ± 0.1 | 0.2 ± 0.0 | 0.1 ± 0.0 | 0.0 ± 0.0 | 0.0 ± 0.0 | 0.1 ± 0.0 |
| 52:2 | 4.2 ± 0.1 | 1.2 ± 0.0 | 0.4 ± 0.0 | 0.3 ± 0.0 | 0.4 ± 0.0 | 0.6 ± 0.0 |
| 52:5 | 2.9 ± 0.1 | 6.6 ± 0.1 | 4.5 ± 0.1 | 5.9 ± 0.4 | 8.4 ± 0.1 | 7.8 ± 0.0 |
| 52:6 | 1.2 ± 0.1 | 9.3 ± 0.2 | 10.5 ± 0.1 | 11.1 ± 0.4 | 12.5 ± 0.1 | 11.1 ± 0.1 |
| 52:7 (isomer 1) | n.d.<sup>c</sup> | 0.0 ± 0.0 | 0.1 ± 0.0 | 0.1 ± 0.0 | 0.1 ± 0.0 | 0.0 ± 0.0 |
| 52:7 (isomer 2) | n.d. | 0.0 ± 0.0 | 0.2 ± 0.0 | 0.4 ± 0.0 | 0.3 ± 0.0 | 0.2 ± 0.0 |
| 52:8 | 0.1 ± 0.1 | 0.0 ± 0.0 | 0.1 ± 0.0 | 0.1 ± 0.0 | 0.6 ± 0.1 | 0.8 ± 0.0 |
| 52:9 | 0.0 ± 0.0 | 0.0 ± 0.0 | 0.2 ± 0.0 | 0.3 ± 0.0 | 0.1 ± 0.0 | 0.0 ± 0.0 |
| 54:0 | 3.0 ± 0.1 | 0.1 ± 0.0 | 0.5 ± 0.1 | 0.4 ± 0.1 | 0.2 ± 0.0 | 0.8 ± 0.0 |
| 54:2 | 0.5 ± 0.0 | 0.1 ± 0.0 | 0.1 ± 0.0 | 0.0 ± 0.0 | 0.1 ± 0.0 | 0.1 ± 0.0 |
| 54:3 | 2.3 ± 0.1 | 0.6 ± 0.0 | 0.3 ± 0.0 | 0.3 ± 0.0 | 0.4 ± 0.0 | 0.5 ± 0.0 |
| 54:4 | 1.6 ± 0.0 | 0.6 ± 0.0 | 0.3 ± 0.0 | 0.4 ± 0.1 | 0.9 ± 0.0 | 1.1 ± 0.0 |
| 54:5 | 2.5 ± 0.1 | 1.8 ± 0.0 | 2.1 ± 0.0 | 3.0 ± 0.2 | 5.0 ± 0.0 | 4.8 ± 0.0 |
| 54:6 | 3.9 ± 0.1 | 4.3 ± 0.0 | 4.4 ± 0.1 | 6.0 ± 0.2 | 7.4 ± 0.0 | 6.9 ± 0.0 |
| 54:7 | 3.1 ± 0.0 | 5.0 ± 0.1 | 2.7 ± 0.0 | 3.6 ± 0.0 | 4.1 ± 0.0 | 3.7 ± 0.0 |
| 54:8 | 3.8 ± 0.1 | 22.0 ± 0.2 | 18.7 ± 0.3 | 21.7 ± 0.3 | 20.8 ± 0.1 | 18.0 ± 0.1 |
| 54:9 | 1.8 ± 0.3 | 41.1 ± 0.4 | 42.8 ± 0.7 | 36.4 ± 0.3 | 32.1 ± 0.1 | 26.9 ± 0.2 |
| 56:0 | 0.2 ± 0.0 | 0.0 ± 0.0 | 0.0 ± 0.0 | 0.0 ± 0.0 | 0.0 ± 0.0 | 0.0 ± 0.0 |
| 56:2 | 0.1 ± 0.0 | 0.0 ± 0.0 | 0.0 ± 0.0 | 0.0 ± 0.0 | 0.0 ± 0.0 | 0.0 ± 0.0 |
| 56:3 | 0.0 ± 0.0 | 0.0 ± 0.0 | 0.0 ± 0.0 | 0.0 ± 0.0 | 0.0 ± 0.0 | 0.0 ± 0.0 |
| 56:4 | 0.0 ± 0.0 | 0.0 ± 0.0 | 0.0 ± 0.0 | 0.0 ± 0.0 | 0.1 ± 0.0 | 0.1 ± 0.0 |
| 56:5 | 0.3 ± 0.0 | 0.0 ± 0.0 | 0.1 ± 0.0 | 0.2 ± 0.0 | 0.6 ± 0.0 | 0.4 ± 0.0 |
| 56:6 | 0.1 ± 0.0 | 0.1 ± 0.0 | 0.6 ± 0.0 | 0.2 ± 0.0 | 0.2 ± 0.0 | 0.2 ± 0.0 |
| 56:7 | 0.0 ± 0.0 | 0.0 ± 0.0 | 0.0 ± 0.0 | 0.0 ± 0.0 | 0.0 ± 0.0 | 0.0 ± 0.0 |
| 56:8 | 0.0 ± 0.0 | 0.0 ± 0.0 | 0.0 ± 0.0 | 0.0 ± 0.0 | 0.0 ± 0.0 | 0.0 ± 0.0 |
| 56:9 | n.d. | 0.0 ± 0.0 | 0.0 ± 0.0 | 0.1 ± 0.0 | 0.0 ± 0.0 | 0.0 ± 0.0 |

<sup>a</sup> The cumulative composition of all three acyl groups are described, with the total number of C atoms represented in the number preceding the colon and the total number of double bonds following the colon.
<sup>b</sup> Percent fraction of total TAG MS1 ion intensity given as mean ± 1 s.e.m, n = 4 biological replicates.
<sup>c</sup> n.d. = not detected

We also looked for evidence of lipid exchange between thylakoids and plastoglobules during the stress. Evidence for the exchange of plastoquinone-9 (PQ-9) was seen, which steadily declined in abundance in thylakoids between 0, 1 and 3 dHS, and reciprocally increased in abundance over the same timeframe in the plastoglobules (**Figure 7B, Supp Tables S10 & S11**). This is suggestive of the movement of PQ-9 from thylakoids to plastoglobules during the heat stress time-course, although we cannot exclude changes in synthesis or degradation as alternate causes. The reason for the subsequent decline of PQ-9 in the plastoglobules from 3 to 7 dHS is unclear but may reflect consumption of PQ-9 due to ROS scavenging or its conversion to various PQ derivatives through hydroxylation and/or acylation (**Supplementary Figures S5**).

The ROS-scavenging compound, α-tocopherol, increased rapidly in the thylakoids over the time course, over 30-fold from 0 to 7 dHS, and remained elevated during the heat recovery. This presumably reflects the role of α-tocopherol in scavenging ROS and quenching lipid peroxidation chain reactions. A somewhat more muted – and delayed - increase (*ca.* 4-fold) was also seen in plastoglobules which remained elevated at 1 dHR but had begun to decrease by 3 dHR (**Figure 7C**). This contrasted with the levels of plastochromenol/plastochromanol (PC-8/PCH_2_-8), also synthesized from Tocopherol Cyclase, whose levels were essentially static in the thylakoid during the time-course and decreased markedly in plastoglobules (**Figure 7C**). Notably, the PC-8 and PCH_2_-8 levels are substantially higher in the plastoglobules than in thylakoids, consistent with their specific plastoglobule localization in *A. thaliana* (Lundquist et al., 2013). In contrast to α-tocopherol, levels of β-carotene steadily and substantially declined in the thylakoid throughout the time-course and declined in the plastoglobules as well, albeit to a smaller extent (**Figure 7B**). The decrease in levels of β-carotene during the stress time-course may reflect turnover of the photosynthetic machinery observed in the proteomic analyses.

As we detailed in our companion paper (Devadasu, et al. 2025 unpublished), a diverse set of annotated plastoquinone derivatives were found in *Z. mays* plastoglobules (and to lesser extent thylakoids) which were hydroxylated on their prenyl tails [PQ(H_2_)-Cs] or acylated on either their head group (PQH_2_-Es) or on the hydroxylation on their side chains [PQ(H_2_)-Bs]. Such compounds were found in plastoglobules and thylakoids from our heat stress study as well and recapitulated the specific use of saturated acyl chains for acylation of PQ-B and PQH_2_-E (**Supplementary Tables S10 & S11**). As for the water-deficit stress treatment (Devadasu, et al., unpublished), cumulative MS1 ion intensity values of each PQ derivative class were among the highest intensities of any lipids (annotated or unannotated) within the plastoglobule dataset, supporting their high levels of accumulation. However, in contrast to the water-deficit stress reported in our companion paper, levels of the PQ derivatives increased during the heat stress time course, typically between 2- and 3-fold (**Supplementary Figures S5 & S6**).

### Identification of fatty acid phytyl esters in *Z. mays* plastoglobules

Fatty acid phytyl esters (FAPEs) are generated by esterification of free phytol with a free fatty acyl group. As free phytol and free fatty acids are both membrane destabilizing, their esterification into FAPEs represents an effective approach to safely store these compounds during rapid turnover of chlorophyll and membrane lipid (Ischebeck et al., 2006; Gaude et al., 2007; Vom Dorp et al., 2015; Spicher et al., 2017; Gutbrod et al., 2019). FAPEs have been shown to accumulate to high levels in *A. thaliana* plastoglobules under N deprivation (which induces chlorophyll turnover and thylakoid disassembly), particularly 16:3-phytol and 12:0-phytol (Gaude et al., 2007). However, the presence of FAPEs in *Z. mays* plastoglobules has not been reported. To determine what FAPEs may accumulate in *Z. mays* plastoglobules, and how their levels may change in response to the heat stress treatment, we isolated and annotated FAPEs using GC-MS. Only 12:0-phytol could be reliably annotated from our dataset (**Figure 8, Supp Fig S7, Supp Table S13**). In contrast to results from *A. thaliana*, we could not find evidence to support annotation of FAPEs comprising any unsaturated acyl groups, including 16:3-phytol. While the 12:0-phytol could be identified in plastoglobules at all time points, a striking 10-fold increase in abundance was seen from 0 to 7 dHS (**Figure 8**). Thus, we conclude that *Z. mays* leaf plastoglobules comprise a 12:0 FAPE species with a short chain saturated acyl group and which is highly induced during the turnover of thylakoids and photosynthetic machinery under our heat stress treatment.

**Figure 8.**
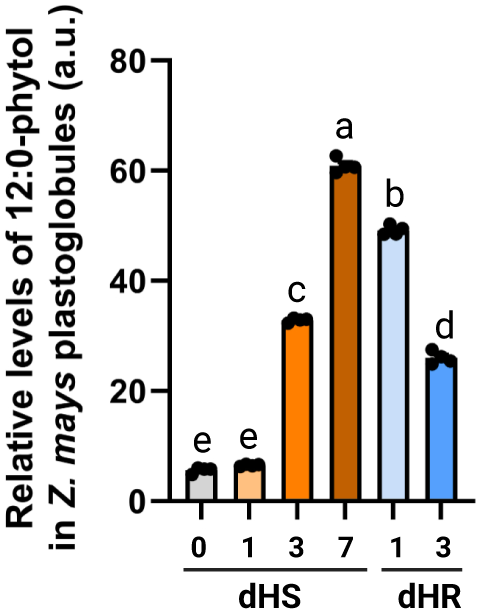
The fatty acid phytyl ester, 12:0-phytol, is found in *Z. mays* plastoglobules and increases in abundance in response to heat stress. Levels are plotted in relative (arbitrary units) at each of the six time points as a bar chart with individual data points as black dots. Error bars indicate mean ± 1 s.e.m. and are not visible when they are smaller than the individual data points. n = 4 biological replicates.

## DISCUSSION

Plastoglobules are ubiquitous throughout the plant lineage suggesting essential roles in supporting photosynthetic life. Yet their functions remain enigmatic. We and others have previously proposed that plastoglobules support membrane remodeling of the thylakoid in response to stress (Lundquist et al., 2012; Rottet et al., 2015; Lundquist et al., 2020; Espinoza-Corral et al., 2021). Such a proposal would suppose an exchange of protein and lipid between the two sub-compartments (Kirchhoff, 2019). Here, we explore this possibility by lipidome and proteome analyses of isolated sub-compartments to establish their dynamic compositions under a heat stress and recovery time-course. Furthermore, prior studies of plant chloroplast plastoglobules have focused on *A. thaliana*. This work represents the first in-depth study of the plastoglobule composition in an agricultural crop. As a monocot, our investigation of *Z. mays* plastoglobules provides an evolutionarily distinct view of plant plastoglobules, highlighting the shared and unique features of the plastoglobule lipidome and proteome.

Notably, TAGs and numerous PQ derivatives [PQ-B, PQ(H_2_)-C, PQH_2_-E] appear to be major constituents of *Z. mays* plastoglobules at all time points. The proportion of total MS1 ion intensity assigned to these species increases significantly under the heat stress, with TAG levels increasing over 10-fold, and PQ-B and PQH_2_-E levels increasing approximately 2-fold, between 0 and 7 dHS. This is in striking contrast to the water-deficit stress where levels decline over 2-fold as a proportion of total MS1 ion intensity. Thus, although TAG and PQ derivatives are major constituents of the *Z. mays* plastoglobules, their relative abundance responds in opposing directions under the two stresses. The TAG may serve as a temporary storage site for acyl groups released from thylakoid lipids during membrane remodeling. Indeed, our lipidomic analyses of the thylakoid demonstrated reduced levels of MGDG and striking depletion of 18:3 acyl groups. In addition, our observed shift in TAG species from more saturated to less saturated acyl groups during the heat stress is consistent with a movement of the 18:3 acyl groups from galactolipids to plastoglobule-localized TAG during heat stress. Correspondingly, TAG species were primarily comprised of 54:9, 54:8 and 52:6, which are consistent with the presence of one or more 18:3 acyl groups.

The function of the PQ derivatives, however, is less clear. Acylation of PQ, as seen in PQ-B and PQH_2_-E species, may serve simply to support their accumulation in the hydrophobic core of the plastoglobules. Nonetheless, PQ(H_2_)-C, and to a lesser extent PQ-B, species are redox-active and capable of transporting electrons in *in vitro* PSII photoreduction reactions (Henninger and Crane, 1966, 1967a, 1967b; Kruk et al., 1998). Hence, a role in photosynthetic regulation cannot be ruled out.

We have also identified a 12:0 FAPE species that accumulates in *Z. mays* plastoglobules. Levels of this FAPE increased 10-fold between 0 and 7 dHS. FAPE species have been identified in *A. thaliana* plastoglobules, primarily in the 12:0 and 16:3 forms, which increase dramatically under N deprivation (Gaude et al., 2007). Importantly, we only found evidence for 12:0 FAPE, not 16:3 FAPE in our *Z. mays* plastoglobules. It was previously suggested that distinct pathways may exist for the synthesis of 12:0 and 16:3 FAPE in *A. thaliana* (Gaude et al., 2007). Our identification of only one of these two FAPE forms is consistent with this notion and indicates that *Z. mays* may only harbor the enzyme(s) for synthesis of the 12:0 FAPE, although we cannot discard the possibility that 16:3-FAPE (or other unsaturated groups) are oxidized during handling rendering them unobservable in our experimental system.

Strikingly, the relative proportions of over half of the plastoglobule proteins were changed during the heat stress. This was despite the generally stable composition of the thylakoid proteome during the heat stress. Among the changes at the plastoglobule proteome, the increased abundance of LOX10/11 and FBA-2a/2c were the most striking. As noted below, these coincided with the appearance of other components of the Calvin-Benson cycle. Surprisingly, the FBN proteins did not increase in proportional abundance, as was seen with FBN1a and FBN1b under the water-deficit stress.

In addition, we observed evidence for the recruitment of multiple enzymes of the Calvin-Benson cycle and JA biosynthesis to the plastoglobule. It has long been recognized that a sub-population of Fructose Bis-phosphate Aldolase (FBA) localizes (apparently constitutively) to *A. thaliana* plastoglobules, although the significance of this localization has remained mysterious (Vidi et al., 2006; Ytterberg et al., 2006). In the present work, we see the same localization of FBA in *Z. mays* and a substantial increase in these and most of the other Calvin-Benson cycle enzymes associated with the isolated plastoglobules. This is even though the levels of these proteins remain unchanged at the total leaf level. No evidence has been reported for the localization of the other Calvin-Benson cycle enzymes at the *A. thaliana* plastoglobules under stressed or unstressed conditions. Thus, the recruitment of these enzymes may be unique to certain species, including *Z. mays*, or unique to certain stresses that have not been tested in the context of *A. thaliana*. Indeed, this recruitment was clearly not seen in our companion paper studying *Z. mays* plastoglobules under water-deficit stress. The physiological significance of the re-mobilization of these Calvin-Benson cycle enzymes is unclear but may represent a way to inactivate the Calvin-Benson cycle under reduced provision of energy and reducing equivalents associated with the heat stress. The significance of the apparent remobilization of LOX10/11 and AOS2b is also unclear but may represent activation of oxylipin biosynthesis – which utilizes the α-linolenic acid that is abundant within these membrane systems – presumably for stimulation of stress signaling and adaptation to the heat stress.

Our results illustrate the dynamic changes to the lipidome and proteome of *Z. mays* plastoglobules and, when placed beside our companion paper investigating response to water-deficiency, highlights the differing responses under the two stresses. Of special note, the elevated levels of TAG and PQ derivatives in the plastoglobules, and recruitment of parts of the Calvin-Benson cycle and JA biosynthesis, distinguish the changes under heat stress from those seen under water-deficit stress. The differing responses presumably reflect the differing physiological manifestations of the stresses which require different adaptive roles for the plastoglobules.

## MATERIALS AND METHODS

### Plant materials, heat treatment and measurement of relative water content

Maize (*Zea mays* ssp. *mays* var. B73) seeds were washed with distilled water twice and kept in the dark overnight. Seeds were then sown in plastic pots (diameter□×□height: 9 ×□10 cm) filled with Sure Mix soil and germinated in a greenhouse with a 16□h photoperiod (natural day light supplemented with light ballasts) and day/night temperatures of 27 and 23□ °C, respectively. Relative humidity was 50–60%. Equal volumes of water (200 mL) were provided to each pot on a weekly basis. Four weeks after germination, when four leaves were fully expanded (*i.e.*, leaf collar is visible) a progressive heat stress was imposed for 7 days. Imposition of stress and recovery was monitored by measuring the relative water content (RWC) of the fifth collared leaf on each day of the time-course approximately two hours after supplemental lights were turned on. Leaf RWC was estimated using the relative turgidity method (Weatherley, 1950). Briefly, the fifth leaf was harvested and immediately measured for fresh weight (FW). Turgid weight (TW) was then determined after leaf segments were immersed in distilled water for 24 h, and finally dry weight (DW) was measured after leaf segments were dried at 65□°C in an oven for 24□h. Relative water content (RWC) was calculated as:

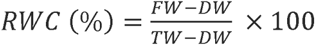

### Collection of tissue and pigment measurements

Aboveground tissue from 8-12 plants, comprising *ca.* 90 g of fresh weight, were pooled together for each biological replicate and used for plastoglobule and thylakoid isolations, chlorophyll measurements, and proteomic and lipidomic analyses. Chlorophyll was measured spectrophotometrically according to Lichtenthaler (1987). Briefly, working in the dark under a green safety lamp at 4 °C, for each replicate 20 mg FW of leaf tissue was ground in liquid N_2_ and mixed with cold 80% (v/v) acetone. The resulting slurry was immediately transferred into a 1.5 mL Eppendorf tube. After incubating with gentle mixing overnight, the samples were centrifuged at 18,000 g for 15 minutes at 4 °C. The supernatant was diluted 10-fold in cold 80% (v/v) acetone and absorbance measured at 663.2, 646.8 and 470.0 nm wavelengths. Concentration estimates were calculated using the following equations:

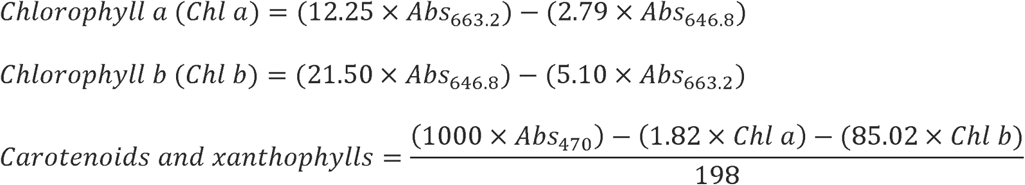

### Photosynthetic measurements

The fifth collared leaf was selected from 13 randomly selected plants at each time point for photosynthetic measurements. Measurements were collected from the middle of the leaf, off the midrib, 2-3 hrs after lights were turned on, using the MultiSpeQ v2.0 device (PhotosynQ Inc., East Lansing, MI, USA) with the RIDES 2.0 parameters, available on the PhotosynQ platform (https://photosynq.org/). Data was plotted for photosynthetic activity (F_v_’/F_m_’), the total energy dissipation as non-photochemical quenching (NPQ_t_), and the quantum yield of photosystem II (□_II_). The photosynthetic measurements were uploaded to the PhotosynQ platform (https://photosynq.org/) with project ID: ‘HT-Peter-2022’ and are publicly available.

### Transmission electron microscopy

The fifth leaf of four randomly selected plants were trimmed at the midpoint of the leaf (halfway between base and tip and off the midrib and sliced into 2 x 2 mm^2^ pieces with a fresh razor blade at the growth chamber before placing on ice for immediate delivery to the laboratory. Tissue sections were then fixed in 2% (w/v) paraformaldehyde and 2 % (w/v) glutaraldehyde in 0.1 M cacodylate buffer by infiltration under vacuum, and incubated overnight at room temperature. The fixed samples were washed in cacodylate buffer and post-fixed in 1% (w/v) osmium tetroxide in the same buffer for 2 h. They were then dehydrated in an ethanol series, transferred to propylene oxide, and embedded in Spurr’s resin. Thin sections (70 nm) were cut with a diamond knife on a Power Tome ultramicrotome (RMC, Boeckler Instruments, Tucson, AZ, USA), and the sections were stained with 2% (w/v) uranyl acetate for 30 min and then with 3% (w/v) lead citrate for 15 min. Observations of chloroplasts were made on a JEM 1400Flash transmission electron microscope (JEOL, Tokyo, Japan) fitted with a high-resolution Tietz F224 digital camera. Morphological measurements from the micrographs were made using ImageJ software available from the web at http://rsbweb.nih.gov/ij/. Transmission electron microscopy of isolated (in vitro) plastoglobules were performed exactly as described in Susanto et al. (2025).

### CD and FT-IR spectroscopy measurements

CD spectra were recorded with a JASCO J-815 (Japan) dichrograph between 400 and 750 nm, with each sample scanned with three accumulations at a scan speed of 100 nm min^−1^. Measurements were carried out at room temperature, with a bandwidth of 4 nm and data pitch of 1 nm. The chlorophyll concentration in thylakoid membrane suspensions was adjusted to 80 μg ml^−1^ and measured in a cuvette with a 1 cm optical path length.

Fourier Transform Infrared (FTIR) spectra were recorded using a Perkin Elmer II 80 V spectrometer equipped with a LiTaO3 detector, with a spectral resolution of 4 cm□¹ and 64 scans per sample. Background single-beam spectra were recorded from an empty ATR plate under a nitrogen purge. Spectra were measured over a range of 800 to 1,800 cm□¹, with protein amide bond I signals detected at 1,657 cm□¹, amide bond II signals at 1,542 cm□¹ and lipid bond signals were detected at 2,925 and 2,854 cm^-1^. An equal dry weight (dw) was used for all samples. Baseline offset corrections were applied to the absorption spectra as needed, using Spectrum 10 software (version 10.5.1, Perkin Elmer, USA).

### Plastoglobule and thylakoid preparations

Plastoglobules were isolated from the aboveground tissue by collecting 12 plants (90 g of tissue) for each biological replicate. Isolation was performed essentially as described in Shivaiah et al. (2022) and was used for isolation of thylakoids and purification of plastoglobules. Briefly, the maize leaves were ground in a Waring blender in 500 mL of grinding buffer (50 mM HEPES-KOH [pH-8.0], 5 mM MgCl_2_, 100 mM Sorbitol] and filtered through gauze. Thylakoid membranes were pelleted by centrifugation of the filtrate at 1500 *g* for 6 min at 4 °C and were resuspended in medium R [[HEPES pH 8.0], 5 mM MgCl_2_, a cocktail of phosphatase and protease inhibitors containing antipain [74 µm] bestatin [130 µm], chymostatin [16.5 µm], E64 [56 µm], leupeptin [2.3 µm], phosphoramidon [37 µm], AEBSF [209 µm], aprotinin [0.5 µm], NaF [50 mM], β-glycerophosphate [25 mM], Na-orthovanadate [1 mM], and Na-pyrophosphate [10 mM]) with 0.2 M sucrose. Several aliquots of the resuspension were flash frozen in liquid N_2_ and stored at -80 °C until use. The remaining thylakoid membranes were aliquoted into four 3.2 mL polycarbonate open-top ultracentrifuge tubes (Beckman Coulter, Pasadena, CA, USA) and tip sonicated with a FB120 sonicator and a 4-tip horn (Fisher Scientific, Hampton, NH, USA). Each tube was pulse sonicated 4 x 10 sec at an amplitude (100%) with a 50 sec pause on ice between each pulse. Sonicated thylakoid suspensions were centrifuged at 150,000 x g for 30 min at 4 °C. The resulting floating pad consisting of crude plastoglobules was collected using a disposable syringe and 22G needle. Crude plastoglobules were then purified on a sucrose gradient of 0.7 M, 0.2 M and 0 M sucrose in medium R. Purified plastoglobules were collected from the top of the gradient, flash frozen, and stored at -80 °C until use.

### Lipid analysis (GC-FID) from thylakoids and plastoglobules

Lipids were extracted as described by Wewer et al. (2013). Briefly, thylakoid samples were adjusted to an equal protein concentration (∼0.1□g) and plastoglobules were adjusted based on OD_700_ levels. Samples were homogenized in 5□mL of chloroform/methanol/formic acid (10:20:1, v/v/v); the homogenate was collected and shaken vigorously. Subsequently, 2.5□mL of 1□M KCl in 0.2□M H_3_PO_4_ was added and the mixture was vortexed briefly. The homogenized samples were centrifuged at 13000*g* for 1 min, and the lower chloroform layer was transferred to a new vial. Extraction was repeated three times by adding 5□mL of chloroform/methanol (2:1, v/v) to the residue, shaking and centrifuging the mixture, and collecting and pooling the chloroform phases. The combined chloroform phases were evaporated with a stream of N_2_ gas, and 100□μL of chloroform was added to dissolve the lipids. Polar lipids were separated with TLC plates treated with (NH4)_2_SO_4_ as described by Wang and Benning (2011). The galacto- and phospho-lipids were marked after iodine staining and scraped off with a razor blade and placed into 1.5 mL Eppendorf tubes. Using pentadecanoic acid (C15:0) as an internal standard, acyl groups were trans-methylated into their fatty acyl methyl esters (FAME) using 1 M HCl in methanol at 80 °C for 20 min. The resulting FAMEs were quantified by a gas-liquid chromatography system (GC-2010; Shimadzu Corp., Kyoto, Japan) equipped with a 30 m DB-23 column with inner diameter of 0.25 mm and film thickness of 0.25 µm (Agilent Technologies, Santa Clara, CA, USA) and with flame ionization detector (FID) according to Zhang et al. (2015).

### Immunoblots

Thylakoids and plastoglobules were separated by SDS-PAGE and solubilized according to Laemmli (Laemmli, 1970). Proteins were transferred on to nitrocellulose membrane using the semi-dry blotting method and probed with primary antibodies (Anti-PsbA, 1:10,000 and anti-FBN1a, 1:5000) purchased from Agrisera AB (product numbers: AS05 084 and AS06 116, respectively; Vännäs, Sweden) followed by incubation with secondary polyclonal anti-rabbit antibody tagged with HRP from Cell Signaling Technology (product number 7074; Danvers, MA, USA) and detected with chemiluminescence using the Amersham ECL kit (product number RPN2232; Chicago, IL, USA) according to the manufacturer’s instructions.

### LC-MS/MS lipid measurements

Total lipids were extracted from freshly isolated thylakoids (based on protein amount of 5 mg) and freshly isolated plastoglobules (OD_700_ = 1.0; 200 µL) with 1 mL of extraction buffer (chloroform:methanol:formic acid (20:10:1 v/v/v)). Samples were shaken vigorously for 5 min, and then 500 µL of water: methanol (3:1 v/v) was added and well mixed. Phases were completely separated by centrifugation for 2 min at 13 000 *g* at room temperature. The upper chloroform phase was removed, and lipids re-extracted three times. Pooled chloroform phases were dried down under a gentle stream of N_2_ gas. Lipids were resuspended in 100 µL isopropanol and diluted 20-fold for LC-MS/MS measurement.

Lipid samples were analyzed using a Waters Xevo G2-XS Q-TOF mass spectrometer interfaced with a Waters Acquity binary solvent manager and Waters 2777c autosampler. Next, 10 µL samples were injected into an Acquity UPLC BEH-C18 column (2.1 x 100 mm) held at 55 °C. The mobile phases consisted of 0.1% (w/v) formic acid and 10 mM ammonium formate in acetonitrile/water (60:40 v/v) (Solvent A) and isopropanol/acetonitrile (90:10, v/v) containing 0.1% (w/v) formic acid and 10 mM ammonium formate (Solvent B). A gradient of mobile phase was applied in a 15-min program with a flow rate of 0.4 mL min ^-^1. The gradient profile was performed as follows: initial conditions were 80% A and 20 % B, ramp to 43% B at 2 min, followed by a linear ramp to 54% B at 12 min, ramp to 70% B at 12.10 min, ramp to 99% B at 18 min, then return to 20% B at 18.1 min and hold until 20 min. LC separated analytes were ionized by both positive and negative ion mode electrospray ionization and mass spectra were acquired using an MS^e^ method in continuum mode over m/z 50 to 1500 to provide data under non-fragmenting and fragmenting conditions (collision energy ramp from 20–80 V). Capillary voltages were set to 3 kV (positive mode) or 2 kV (negative mode), cone voltage was 30 V, source temperature was 100°C, desolvation temperature was 350°C, desolvation gas flow was 600 L/hr and cone gas flow was 40 L/hr. Data were processed using Masslynx and Progenesis QI software (Waters Corp, Milford, MA, USA).

### Proteomics sample preparation

Proteomic analysis of thylakoids and plastoglobules was performed with lyophilized thylakoids normalized on equal amount of protein concentration (100 µg) and lyophilized plastoglobules normalized to 1.0 OD_700_. Both thylakoids and plastoglobules were re-suspended in Laemmli sample buffer and heated at 60 °C for 10 min. Samples were cooled and loaded onto a 12.5% pre-cast Criterion 1D gel (Bio-Rad, Hercules, CA, USA) and electrophoresed at 50 V until the Coomassie dye front migrated 2-3 mm below the well. The gel was stained using Coomassie Brilliant Blue G250 (0.05% (w/v) in 50% (v/v) methanol, 10% (v/v) glacial acetic acid and 40% (v/v) water. Concentrated sample bands were then excised from the gel and placed into individual microfuge tubes.

Gel bands were digested in-gel according to Shevchenko, et al. (1996) with modifications. Briefly, gel bands were dehydrated using 100% acetonitrile and incubated with 10 mM dithiothreitol in 100 mM ammonium bicarbonate, pH∼8, at 56 °C for 45 min, dehydrated again and incubated in the dark with 50 mM chloroacetamide in 100 mM ammonium bicarbonate for 20 min. Gel bands were then washed with ammonium bicarbonate and dehydrated again. Sequencing-grade modified trypsin was prepared to 0.005 µg/µL in 50 mM ammonium bicarbonate and ca. 100 µL of this was added to each gel band so that the gel was completely submerged. Bands were then incubated at 37 °C overnight. Peptides were extracted from the gel by water bath sonication in a solution of 60 % (v/v) acetonitrile (ACN) / 1% (v/v) trifluoroacetic acid (TFA) and vacuum dried to *ca.* 2 µL.

### LC-MS/MS proteomics

The peptides were resuspended in 20 µL of a solution of 2% (v/v) acetonitrile, 0.1% (v/v) triflouroacetic acid and 97.9% (v/v) water. A sample injection of 5 µL was made using a Thermo (www.thermo.com) EASYnLC 1200 onto a Thermo Acclaim PepMap RSLC 0.075mm x 250mm C18 column and washed for *ca.* 5 min with buffer A (99.9% water / 0.1% formic acid; v/v). For thylakoid samples, bound peptides were then eluted over 45 min with a linear gradient of 8% buffer B (80% acetonitrile / 0.1% formic acid / 19.9% water; v/v/v) to 35% buffer B in 34 min at a constant flow rate of 300 nL min^-1^. After the gradient the column was washed with 90% buffer B for the duration of the run. For plastoglobule samples, bound peptides were eluted over 35 min with a linear gradient of 8% buffer B to 35% buffer B in 24 min at a constant flow rate of 300 nL min^1^. After the gradient the column was washed at 90% buffer B for the duration of the run. Column temperature was maintained at a constant temperature of 50 °C using and integrated column oven (PRSO-V2, Sonation GmbH, Biberach, Germany). Eluted peptides were sprayed into a ThermoScientific Q-Exactive HF-X mass spectrometer (www.thermo.com) using a FlexSpray spray ion source. Survey scans were taken in the Orbitrap (60000 resolution, determined at m/z 200) and the top 15 ions in each survey scan were subjected to automatic higher energy collision induced dissociation with fragment spectra acquired at a resolution of 15000.

### LC-MS proteomic data analysis

The resulting MS/MS spectra were converted to peak lists using MaxQuant v2.4.5.0 (Cox and Mann, 2008) and searched against the *Z. mays* (v3) full peptide dataset (63241 unique protein sequences downloaded from MaizeGDB [https://www.maizegdb.org/]), and concatenated with common laboratory contaminants using the Andromeda search algorithm (Cox et al., 2011). Oxidation of methionine, phosphorylation of threonine, serine, and tryptophan, and N-terminal acetylation were set as variable modifications, carbamidomethylation was set as a fixed modification. Digestion mode was Trypsin/P with a maximum of two missed cleavages. Label-free quantification was enabled with default settings. MS/MS tolerance of the first search was 20 ppm, and the main search was 4.5 ppm, with individualized peptide mass tolerance selected. FDR at peptide spectrum match and protein levels was set as 0.01. Plastoglobules samples and thylakoid samples were searched in separated groups of 0.7 min match time window and 20 min alignment window for match between runs. The ‘primary gene model’ for each protein group was identified as having the highest number of Unique + Razor peptide counts when compiled across all experiments. In the case of a tie, the lowest gene model number was selected, regardless of whether the protein group contained gene models from one or more gene loci. The mass spectrometry proteomics data have been deposited to the ProteomeXchange Consortium via the PRIDE (Perez-Riverol et al., 2022) partner repository (https://www.ebi.ac.uk/pride/) in MIAPE-compliant format with the dataset identifiers PXD061470, PXD061472 and PXD061473.

### Fatty acid phytyl ester measurements

Fatty acid phytyl esters (FAPEs) were identified and quantified from isolated plastoglobules samples. The lipid ester fraction was isolated from being extracted with chloroform/methanol (2:1), and the organic solvent evaporated with nitrogen gas. Lipid esters were dissolved in hexane and injected directly into GC–MS. GC-MS was performed using an Agilent 7890A GC / single quadrupole mass spectrometer with 5975C inert XL MSD (Agilent, Santa Clara, CA). One µL of the sample was injected in a splitless mode with an injector temperature of 300 °C, a purge flow of 40 mL/min at 1 min, and a flow rate of 1.0 mL/min helium. Separation was achieved on an Agilent J&W VF5-ms column (30 m x 0.25 mm x 0.25 mm) (Agilent, Santa Clara, CA) using the following temperature profile: 80°C for 1 min; 40°C min^-1^ to 275°C; 10°C min^-1^ to 325°C; 320°C for 15 min. Ionization employed 70 eV electron ionization and the mass spectrometer was operated in selected ion mode (SIM). Three fragments ions, m/z 123, 124, and 278 were used to detect phytyl dodecanoate and phytyl hexadecanoate. The dwell time for all ions was 100 ms. Phytyl dodecanoate was quantified using a phytyl hexadecanoate standard (synthesized by Dr. A. Daniel Jones III).

### Graphing and statistical analyses

Statistical analyses and plotting were performed with Prism version 9.3 (GraphPad, Boston, MA, USA).

## ACKNOWLEDGEMENTS & FUNDING

This work is supported by USDA National Institute of Food and Agriculture [grant no. 2021-67013-33756] to P.K.L. We thank Dr. John Froehlich for providing the PsbS and Histone3 antibodies. The authors wish to thank Doug Whitten of the MSU Proteomics Core Facility for assistance with proteomic analysis, and Alicia Withrow of the Center for Advanced Microscopy for assistance with TEM.

## AUTHOR CONTRIBUTION

P.K.L conceptualization; P.K.L data curation; E.D., F.S., A.L.S., C.J., P.K.L.; formal analysis; P.K.L funding acquisition; E.D., F.S. investigation; E.D., F.S., A.L.S., C.J., P.K.L. methodology; P.K.L. project administration; P.K.L. supervision; E.D., P.K.L. visualization; E.D., P.K.L. writing – original draft; E.D., F.S., A.L.S., C.J., P.K.L writing – review & editing.

## SUPPLEMENTARY MATERIALS

### Supplementary Figures

**Supplementary Figure S1.** Characterization of the heat-stress treatment on maize.

**Supplementary Figure S2.** Transmission electron micrograph of a bundle sheath cell at 1 dHR.

**Supplementary Figure S3.** Quantitative proteomics of heat shock proteins and selected sub-cellular compartments from total leaf protein samples.

**Supplementary Figure S4.** Quantitative proteomics of selected Calvin-Benson cycle enzymes from total leaf, thylakoids, and plastoglobules during the heat stress time course.

**Supplementary Figure S5.** Relative levels of plastoquinone derivatives in *Z. mays* plastoglobules and thylakoids.

**Supplementary Figure S6.** GC-MS provides experimental support for annotation of 12:0-phytol.

### Supplementary Tables

**Supplementary Table S1.** Leaf relative water content at each time point of the heat stress treatment

**Supplementary Table S2.** Photosynthetic measurements at each time point

**Supplementary Table S3.** Ultrastructure measurements of *Z. mays* under heat stress

**Supplementary Table S4.** Total leaf proteomics analysis

**Supplementary Table S5.** Quantitative proteome of isolated thylakoids during heat stress

**Supplemental Table S6.** All protein identifications from isolated plastoglobules

**Supplementary Table S7.** Quantitative plastoglobule proteome during a heat stress treatment

**Supplementary Table S8.** Polar lipid ratios of isolated thylakoids

**Supplementary Table S9.** Acyl chain composition of polar lipids of thylakoids

**Supplementary Table S10.** Thylakoid LC-MS/MS lipidome of *Z. mays* leaf tissue

**Supplementary Table S11.** Plastoglobule LC-MS/MS lipidome from *Z. mays* leaf tissue

**Supplemental Table S12.** Fatty acid phytyl ester annotations from isolated *Z. mays* plastoglobules

