## Supplementary Figures for "Dynamic changes to the plastoglobule lipidome and proteome in heat-stressed maize"

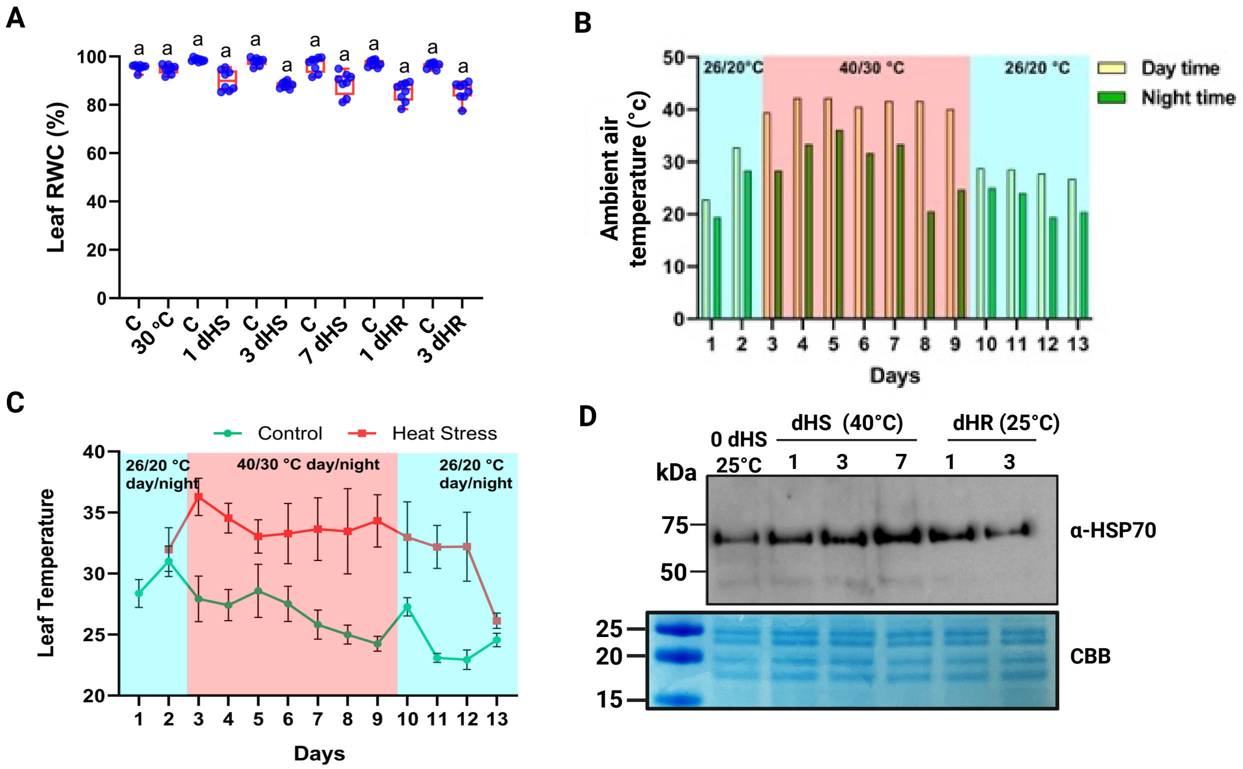


**Supplementary Figure S1**. Characterization of the heat-stress treatment on maize. **A)** Leaf relative water content (RWC) of the fourth collared leaf during the heat treatment at specified time points; n = 8 biological replicates (*i.e.*, individual plants). Box and whisker plots represent the 25th and 75th quartiles with individual data points overlaid as blue circles. **B**) Greenhouse ambient air temperature recordings taken at mid-day of the respective time points during the stress time-course. **C)** Leaf temperatures of the fifth collared leaf, measured with a hand-held MultiSpeQ device, at respective time points during the stress time-course. **D)** Immunoblot of total leaf protein samples against the Heat Shock Protein 70, a marker protein for heat stress in maize chloroplasts.

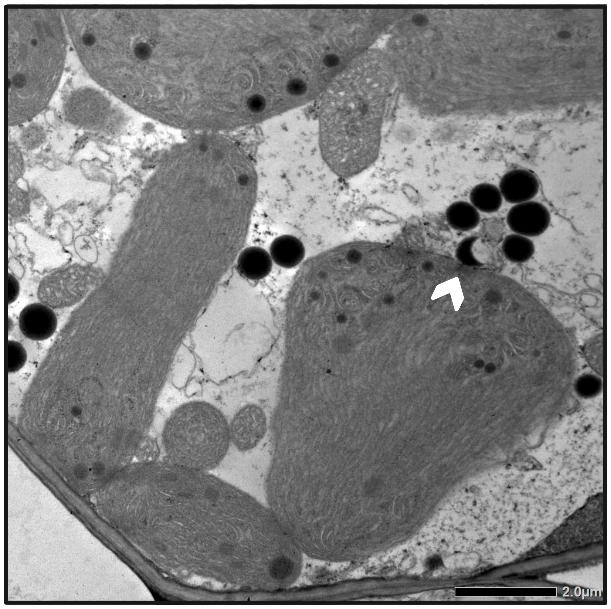


**Supplementary Figure S2**. Transmission electron micrograph of a bundle sheath cell at 1 dHR. Note the prevalent cytosolic lipid droplets and crescent staining pattern among the lipid droplet marked by a white arrowhead. The scale bar designates 2 μm.

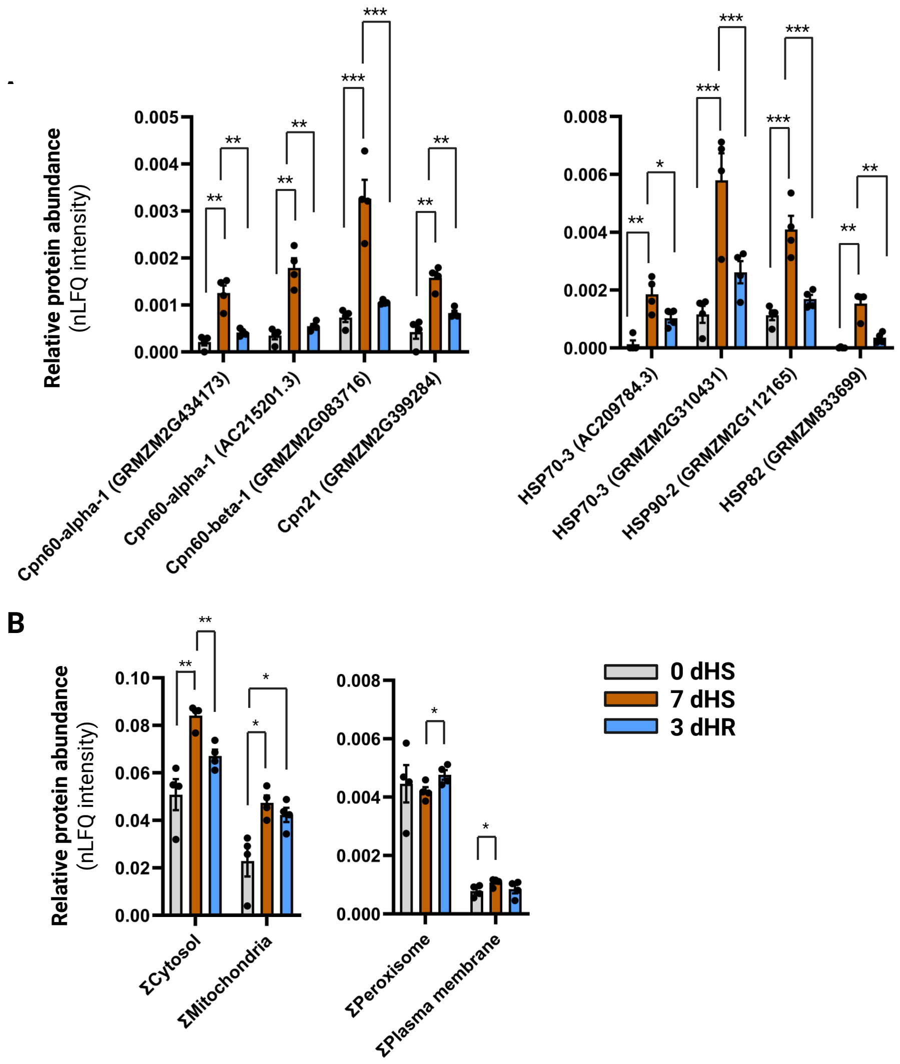


**Supplementary Figure S3**. Quantitative proteomics of heat shock proteins and selected sub-cellular compartments from total leaf protein samples. **A**, relative protein abundance of selected heat shock proteins. Gene locus is indicated in brackets. **B**, nLFQ intensities of all proteins annotated with a localization in the respective sub-compartment according to the Plant Proteome Database were summed. Data are represented as bar graphs plotting the mean ± 1 s.e.m, n = 4 biological replicates (*i.e.*, individual plants). Individual data points are indicated as black dots. * $p < 0.05$, ** $p < 0.01$, *** $p < 0.001$, homoscedastic two-tailed Student's t-test.

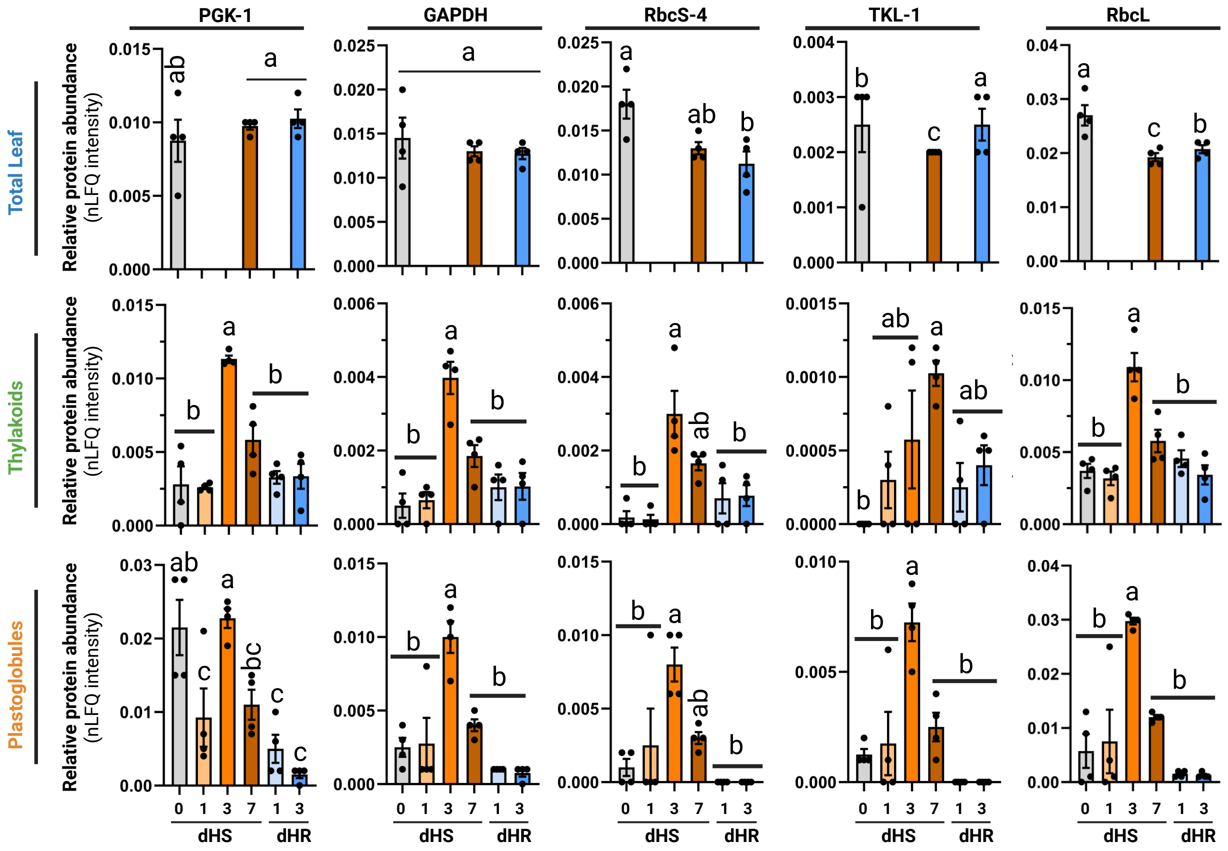


**Supplementary Figure S4.** Quantitative proteomics of selected Calvin-Benson cycle enzymes from total leaf, thylakoids, and plastoglobules during the heat stress time course. Relative protein abundance is plotted as the mean ± 1 s.e.m., n = 4 biological replicates (*i.e.*, individual plants). In addition to the Fructose bis-phosphate 2 isoforms, these four Calvin-Benson cycle enzymes (Phosphoglycerate kinase 1 [PGK-1], Glyceraldehyde-3-phosphate Dehydrogenase 4 [GAPDH], Rubisco small subunit 4 [RbcS-4], and Transketolase 1 [TKL-1]) appear to be remobilized to the thylakoids and/or plastoglobules during the heat stress, as their levels are substantially elevated in the thylakoid and plastoglobule samples, but not at the total leaf level. Note that the total leaf proteome was only assessed at three of the time points. Also note that GAPDH and TKL-1 represent protein groups for which only the majority protein identification is specified in the headers. Statistical significance of differences was assessed using ordinary one-way ANOVA with $p < 0.001$, indicated by lowercase letters. For more information, refer to Supplementary Tables S4-S7.

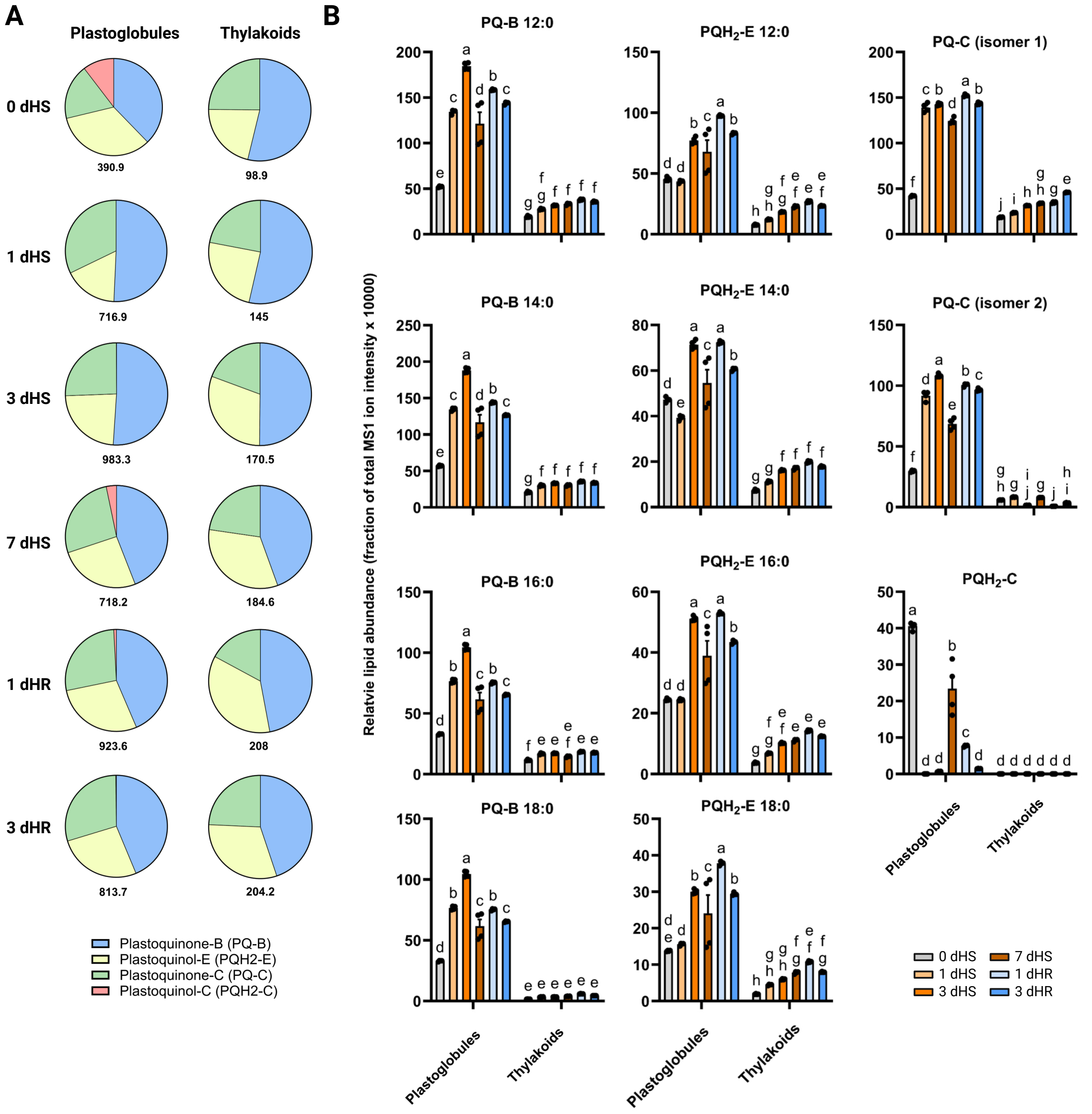


**Supplementary Figure S5.** Relative levels of plastoquinone derivatives in *Z. mays* plastoglobules and thylakoids. **A**, Pie chart of mean cumulative MS1 ion intensities of each class of PQ derivatives in isolated plastoglobules and isolated thylakoids at each time point, n = 4 biological replicates. The number below each pie indicates the sum total of MS1 ion intensity comprised of the four PQ derivative classes. **B**, Bar graphs depicting relative levels of each individual PQ derivative class identified in plastoglobules and thylakoids. Mean ± 1 s.e.m with black dots representing individual data points, n = 4 biological replicates (*i.e.*, individual plants). Statistical significance of differences was assessed using ordinary two-way ANOVA with $p < 0.001$ and Fischer's LSD, indicated by lowercase letters.

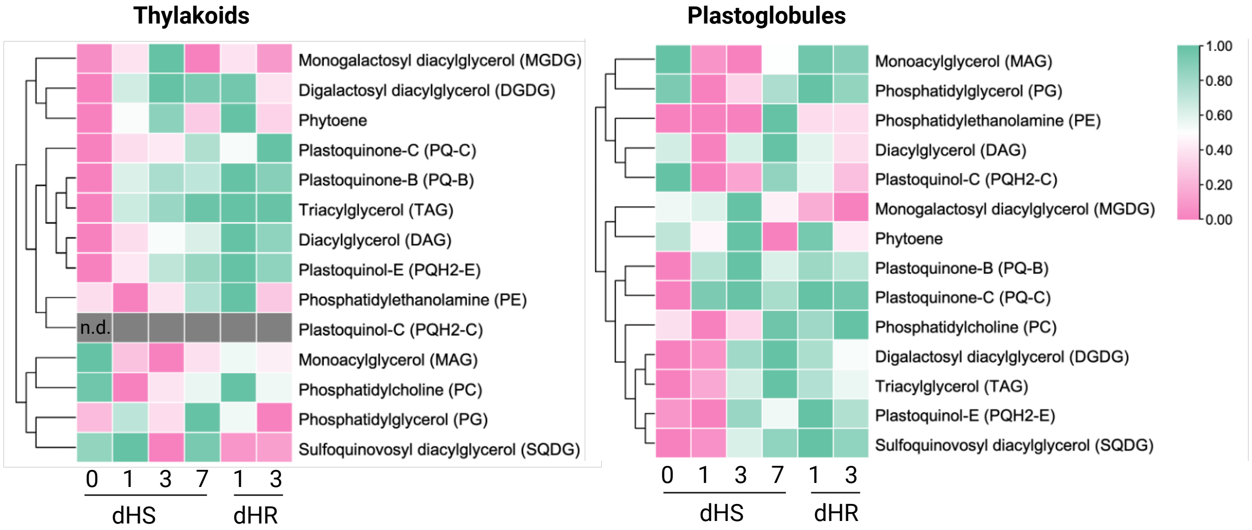


**Supplementary Figure S6.** Relative levels of polar lipids and triacylglycerols in *Z. mays* plastoglobules and thylakoids. Heat map of log2 MS1 ion intensities of each class of polar lipids, TAGs and PQ derivatives in isolated thylakoids and plastoglobules at each time point, n = 4 biological replicates (*i.e.*, individual plants).For more information, refer to Supplementary Tables S10 and S11. *n.d.* = not detected.

**12:0-phytol**

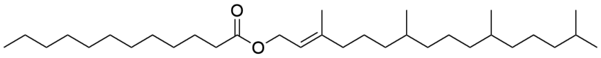


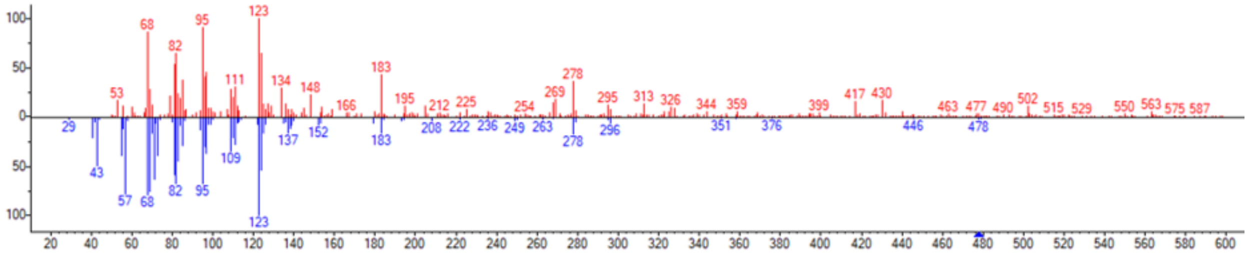


**Supplementary Figure S7.** GC-MS provides experimental support for annotation of 12:0-phytol. The experimental MS spectra is provided above in red. Below, in blue, is the theoretical MS spectra of 12:0-phytol.
